# Coordination of transcription-coupled repair and repair-independent release of stalled RNA polymerase II in response to transcription-blocking lesions

**DOI:** 10.1101/2024.07.07.602436

**Authors:** Yongchang Zhu, Xiping Zhang, Meng Gao, Yanchao Huang, Yuanqing Tan, Avital Parnas, Sizhong Wu, Delin Zhan, Sheera Adar, Jinchuan Hu

**Affiliations:** Shanghai Fifth People’s Hospital, Fudan University, and Shanghai Key Laboratory of Medical Epigenetics, International Co-laboratory of Medical Epigenetics and Metabolism (Ministry of Science and Technology), Institutes of Biomedical Sciences, Fudan University, Shanghai 200032, China; Department of Microbiology and Molecular Genetics, The Institute for Medical Research Israel-Canada, The Faculty of Medicine, The Hebrew University of Jerusalem, Ein Kerem, 9112102, Jerusalem, Israel; These authors contributed equally: Yongchang Zhu, Xiping Zhang, Meng Gao

## Abstract

Transcription-blocking lesions (TBLs) stall elongating RNA polymerase II (PolII), which then initiates transcription-coupled repair (TCR) to remove TBLs and allow transcription recovery. In the absence of TCR, eviction of lesion-stalled PolII is required for alternative pathways to address the damage, but the mechanism is unclear. This study, utilizing Protein-Associated DNA Damage Sequencing (PADD-seq), reveals that the p97-proteasome pathway can evict lesion-stalled PolII independently of repair. Both TCR and repair-independent eviction require CSA and ubiquitination. However, p97 is dispensable for TCR and PolII eviction in TCR-proficient cells, highlighting repair’s prioritization over repair-independent eviction. Moreover, ubiquitination of RPB1-K1268 is important for both pathways, with USP7’s deubiquitinase activity promoting TCR without abolishing repair-independent PolII release. In summary, this study elucidates the fate of lesion-stalled PolII, and may shed light on the molecular basis of genetic diseases caused by the defects of TCR genes.

## Introduction

Successful transcription plays a crucial role in cellular activities^1^. However, elongating RNA polymerases stall at transcription-blocking lesions (TBLs) induced by factors such as UV, cisplatin and benzopyrene^2,3^. Transcription coupled repair (TCR), a sub-pathway of nucleotide excision repair (NER), is the major mechanism to remove TBLs and recover transcription^4,5^. In mammalian cells, TCR is initiated by lesion-stalled RNA polymerase II (PolII) which recruits the Cockayne syndrome group B protein (CSB) to translocate PolII and overcome non-blocking lesions like 8-oxo-deoxyguanosine^6–8^. If the damage is too bulky to be bypassed (i.e., TBLs), Cockayne syndrome group A protein (CSA) is recruited to damage sites in complex with the damage-specific DNA binding protein 1 (DDB1), cullin 4A (Cul4A), ring box 1 (Rbx1), and ubiquitin E3 ligase (CRL4^CSA^)^9,10^. This is followed by the recruitment of the complex of UV-stimulated scaffold protein A (UVSSA) and ubiquitin-specific protease 7 (USP7) ^11–13^. CRL4^CSA^ ubiquitylates surrounding proteins including PolII, CSB and UVSSA^14–16^, while UVSSA-USP7 is thought to protect them from degradation by its deubiquitinase activity^11–13^, thus promoting repair. More importantly, UVSSA interacts with the p62 subunit of transcription factor IIH (TFIIH) to help recruit the core repair factor TFIIH^17^. TFIIH contains two helicases, namely xeroderma pigmentosum complementation group proteins B and D (XPB and XPD), and is also the scaffold for downstream repair factors replication protein A (RPA) and xeroderma pigmentosum complementation group proteins A, F and G (XPA, XPF and XPG)^17,18^. The two endonucleases XPF and XPG incise the damaged strand and release an oligonucleotide containing the lesion, resulting in a gap filled by DNA polymerases and ligases^19,20^.

As an alternative to TCR, the adducts can be removed by the global genome repair (GGR) sub-pathway of NER, or bypassed by translesion DNA synthesis during replication^21,22^. When TCR is blocked, the PolII-damage complex may be even more harmful than the damage itself, since lesion-stalled PolII can hinder the access of these alternative mechanisms to the damage^23,24^. In-vitro, this PolII-damage complex is quite stable^25^, thus lesion-stalled PolII needs to be actively removed if TCR cannot be completed^26^. It was thought that UV-induced ubiquitination of PolII plays an important role in this process^15,27^. Ubiquitinated proteins can be recognized by the ubiquitin-selective protein segregase valosin-containing protein (VCP)/p97 and directed to the 26S proteasome where they undergo proteolysis^28^. Indeed, p97 has been implicated in the proteasome-medicated eviction of RNA PolII^29,30^. Of note, since lesion-stalled PolII directly engages in the recruitment of TFIIH^17^, a pathway that releases PolII without repair could compete with TCR. These two processes must therefore be coordinated to ensure efficient TCR while preventing the accumulation of lesion-stalled PolII. However, the mechanism of this eviction and its coordination with repair remains to be elucidated.

A major challenge for studying the above questions is how to measure the interaction between PolII and DNA damage, as both of them are widely spread across the genome. Furthermore, in addition to stalling PolII in cis, UV-induced lesions have trans-effects on PolII, including reduced levels of initiating Pol II^31^ and slower elongating rates^32,33^, which can alter measurement of PolII in chromatin, making it difficult to assess the direct interaction between PolII and DNA damage^34^. Recently we developed Protein-Associated DNA Damage Sequencing (PADD-seq) to check direct protein-DNA damage interactions by measuring damage distribution in protein-bound DNA fragments obtained from chromatin immunoprecipitation (ChIP) (**Fig. 1a**)^35^. Based on previous PADD-seq measurements, interactions between PolII and UV-induced cyclobutane pyrimidine dimers (CPDs) disappear within two hours in TCR-proficient cells, while PolII was retained at damage sites in *CSB* knockout cells^35^. In this study, focusing on CRL4^CSA^ ubiquitin E3 ligase, the p97-proteasome pathway, the ubiquitination site of PolII and the UVSSA-USP7 deubiquitinase complex, we systematically investigated the roles of ubiquitination and deubiquitylation in both TCR and repair-independent PolII release to unveil the coordination of these two pathways in response to TBLs.

**Fig. 1.**
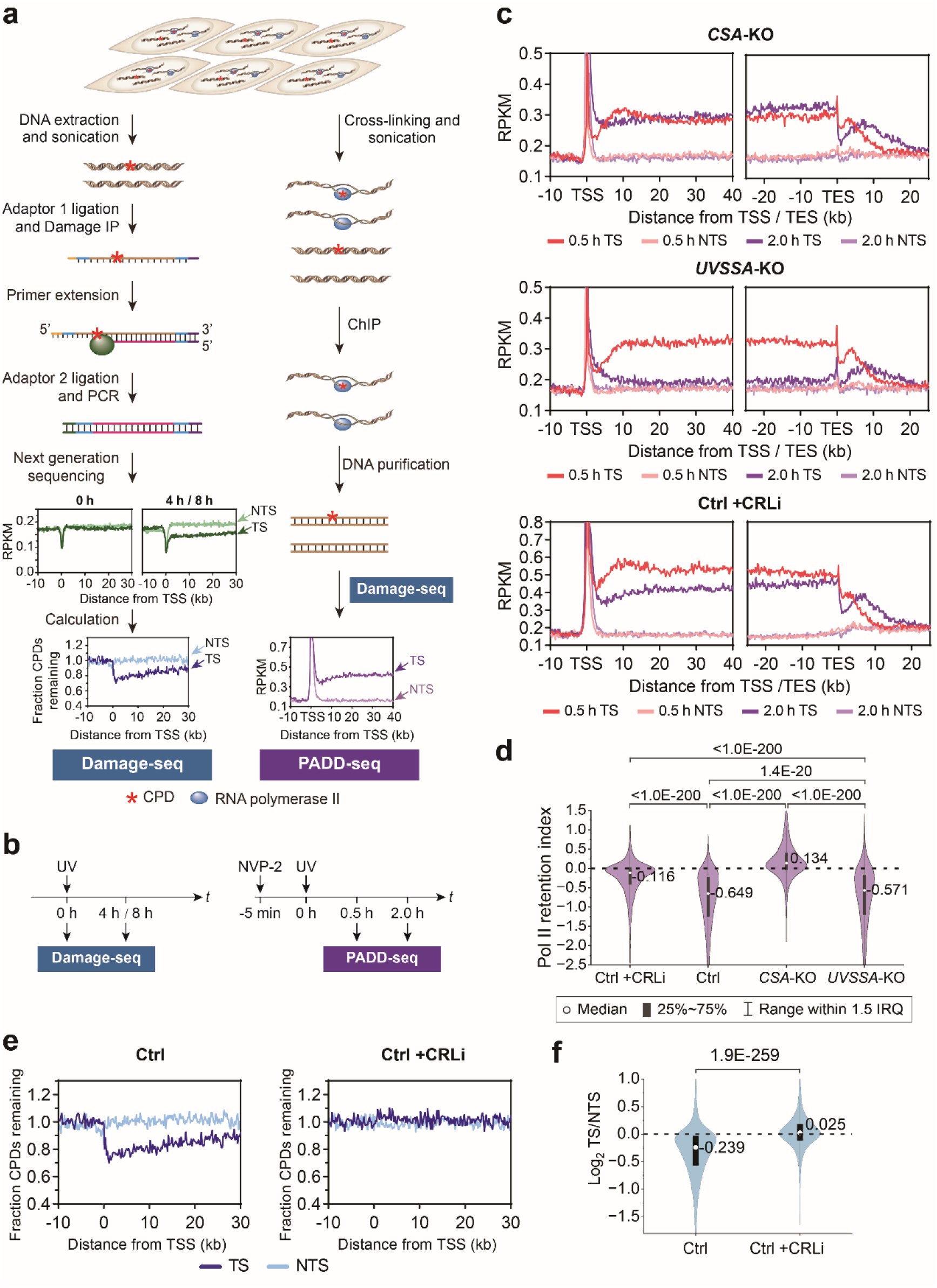
Lesion-stalled PolII is resolved in *UVSSA*-KO cells by a CSA- and ubiquitination- dependent manner. **a,** Schematic representation of Damage-seq and PADD-seq. For Damage-seq, genomic DNA extracted from UV-irradiated cells are sonicated and ligated to the first adaptor, followed by immunoprecipitation with the lesion-specific antibody. The precise sites of the lesions are determined by the stalling of primer extension with high-fidelity DNA polymerase. After the ligation of the second adaptor, the primer extension products are PCR-amplified and sequenced. For PADD-seq, PolII-associated DNA are enriched by regular chromatin- immunoprecipitation (ChIP) with an anti-pan-RPB1 antibody. Purified DNA fragments are subjected to Damage-seq. **b,** Diagram of the experimental design for the measurement of TCR (by Damage-seq) and PolII-CPD interaction (by PADD-seq). **c,** Meta-gene analysis of PADD-seq signals around TSSs and TESs for active genes longer than 50 kb (*n* = 2790) under indicated conditions. **d,** Quantification of PolII retention on damage sites by relative change of PADD-seq signals from 0.5 h to 2 h on each gene. Active genes longer than 20 kb were selected (*n* = 4488). **e,** Meta-gene analysis of Damage-seq signals around TSSs for active genes longer than 50 kb (*n* = 2790) under indicated conditions. Cells were collected immediately (0 h) or at 8 h after UV irradiation. Fraction CPDs remaining was calculated as the ratio of 8 h to 0 h. **f,** Violin plots of relative Damage-seq signals on each active gene (*n* = 6406). Log2 value of the ratio of fraction CPDs remaining on TS to that on NTS was calculated. *P* value was calculated using two-tailed paired Student’s *t*-test for **(d)** and **(f)**. See also Supplementary Fig. 1 and 2. Source data are provided as a Source Data file.

## Results

### Lesion-stalled PolII is resolved in *UVSSA*-KO cells by a CSA- and ubiquitination-dependent manner

To determine the fate of lesion-stalled PolII, we took advantage of PADD-seq to measure PolII-CPD interaction in an XPC-deficient cell line (XP4PA-SV-EB, henceforth referred to as XP-C cells) that lacks GGR while possessing proficient TCR to avoid the interference of GGR^36^. To avoid re-stalling of the next PolII after eviction of a stalled PolII, cells were treated with a CDK9 inhibitor (NVP-2) prior to UV to prevent *de novo* PolII promoter-release after UV treatment (**Fig. 1b**, **right**)^35^. Under this condition, there is always a sharp peak of PADD-seq at transcription start sites (TSSs) (**Fig. 1c**) due to high PolII occupancy, although there are fewer CPDs around TSSs owing to high GC content in this region^37,38^. Moreover, the downstream regions of transcription end sites (TESs) have high density of slow-moving PolII^39^, resulting in persistent PADD-seq peaks which have a shift towards the downstream direction from 0.5 h to 2 h (**Fig. 1c**)^35^. In this study we focused on the fate of lesion-stalled elongating PolII, thus only PADD-seq signals in the gene body region (from TSS downstream 10 kb to TES, see Methods) were taken into account. Since UV induces CPDs on both template strands (TSs) and non-template strands (NTSs) independent of PolII occupancy^37^, the signals of PolII-associated CPDs detected by PADD-seq should be weak on both strands right after UV irradiation. After a certain time (0.5 h in our experiments), elongating PolII would be blocked by lesions on TSs but not those on NTSs, thus specific PADD-seq signals on TSs reflecting the level of lesion-stalled PolII could be detected. Then since under these experimental conditions, *de novo* PolII release is inhibited, PADD-seq signals would decrease and finally disappear if pre-existing PolII blocked by the lesions can be resolved (**Supplementary** Fig. 1a). Our previous data showed that PADD-seq signals disappeared at 2 h after UV treatment in the presence of a CDK9 inhibitor in TCR-proficient XP-C cells but not in *CSB*-KO cells^35^.

To explore the roles of CSA and UVSSA in determining the fate of lesion-stalled PolII, *CSA*-KO and *UVSSA*-KO XP-C cells were generated by CRISPR-Cas9 and verified by Sanger sequencing and Western blot (**Supplementary** Fig. 1b-e). PADD- seq was performed in *CSA*-KO and *UVSSA*-KO cells to assess the retention of PolII on CPDs at 0.5 and 2 hours after UV treatment (**Fig. 1b**, **right**). As previously reported, PADD-seq could detect lesion-stalled PolII at 0.5 h after UV irradiation under all tested conditions^35^. Similar to *CSB*-KO cells^35^, PolII was restrained at damage sites for 2 hours in *CSA*-KO cells (**Fig. 1c top and Supplementary** Fig. 2a **top**), suggesting that lesion-stalled PolII could not be removed in the absence of either CSB or CSA. In sharp contrast, PolII was efficiently released from CPDs in *UVSSA*-KO cells within 2 hours (**Fig. 1c middle; Supplementary** Fig. 2a **middle and 2b top**). Although loss of UVSSA can cause complete inactivation of TCR^11–13,17^, the dynamics of PolII-CPD interaction in *UVSSA*-KO cells is similar to TCR-proficient XP-C (Ctrl) cells (**Fig. 1d**; **Supplementary** Fig. 2b **bottom and 2c**), suggesting a repair-independent but CSA-dependent pathway to release lesion-stalled PolII. Since CRL4^CSA^ is a cullin family ubiquitin E3 ligase^40^, it is speculated that UV-induced ubiquitination of PolII and other TCR factors may play a role in this repair-independent process^14,41,42^. Therefore, we tested the role of cullin-dependent ubiquitination and found that the cullin family E3 ligases inhibitor (CRLi)^43^ suppressed PolII release in TCR-proficient cells (**Fig. 1c bottom and 1d; Supplementary** Fig. 2a **bottom and 2c**). We also checked the effect of cullin-dependent ubiquitination on TCR by Damage-seq which can map the genomic distribution of lesions at base resolution in a strand-specific manner. Since TCR selectively removes lesions on TSs, there should be less damage on TSs than on NTSs in TCR-proficient cells after incubation that allows repair (**Fig. 1a,b**, **left**). As shown in **Fig1. e,f,** CRLi could also abrogate TCR in XP-C cells, indicating an essential role of ubiquitination in TCR. Therefore, cullin-dependent ubiquitination is indispensable in both TCR and repair-independent PolII release. Moreover, UVSSA-KO cells also showed stronger eviction of stalled PolII upon cisplatin treatment compared to CSA- KO cells, suggesting the versatility of repair-independent PolII release mechanism across different TBLs (**Supplementary** Fig. 2d,e). However, TCR-proficient XP-C cells displayed only weak accumulation of lesion-stalled PolII, likely due to concurrent damage formation and repair during cisplatin treatment (**Supplementary** Fig. 2d).

Subsequently, we examined the UV-induced PolII ubiquitination in *CSA*-KO and *UVSSA*-KO cells. Intriguingly, loss of CSA could not completely abolish UV-induced PolII poly-ubiquitination (henceforth referred to as PolII ubiquitination) (**Supplementary** Fig. 2f), as previously reported^15,27^. However, our results suggested that CSA-independent PolII ubiquitination could not support either TCR or repair-independent PolII release, implying that lesion-stalled PolII could only be ubiquitinated in a CSA-dependent manner. In contrast, although loss of UVSSA also reduced UV-induced PolII ubiquitination as previously reported^15^ (**Supplementary** Fig. 2f), lesion-stalled PolII is efficiently removed under this condition.

### p97 evicts PolII from damage sites in the absence of UVSSA, but is dispensable for both TCR and PolII release in TCR-proficient cells

Since repair-independent PolII release required cullin-dependent ubiquitination, we asked whether it is driven by p97 and the proteasome. PADD-seq revealed that the PolII-CPD interaction on TSs persists in *UVSSA*-KO cells in the presence of either a p97 inhibitor or a proteasome inhibitor (**Fig. 2a,b and Supplementary** Fig. 3a-c), indicating that lesion-stalled PolII is evicted by the p97-proteasome pathway in the absence of UVSSA. Thus, although UV-induced PolII ubiquitination is reduced in *UVSSA*-KO cells, the remaining ubiquitination on lesion-stalled PolII is sufficient to support p97-mediated release. Furthermore, the p97 inhibitor also prevented PolII release from cisplatin-damage sites in *UVSSA*-KO cells (**Supplementary** Fig. 3d,e), implying that p97 could eliminate PolII blocked by a wide range of TBLs.

**Fig. 2.**
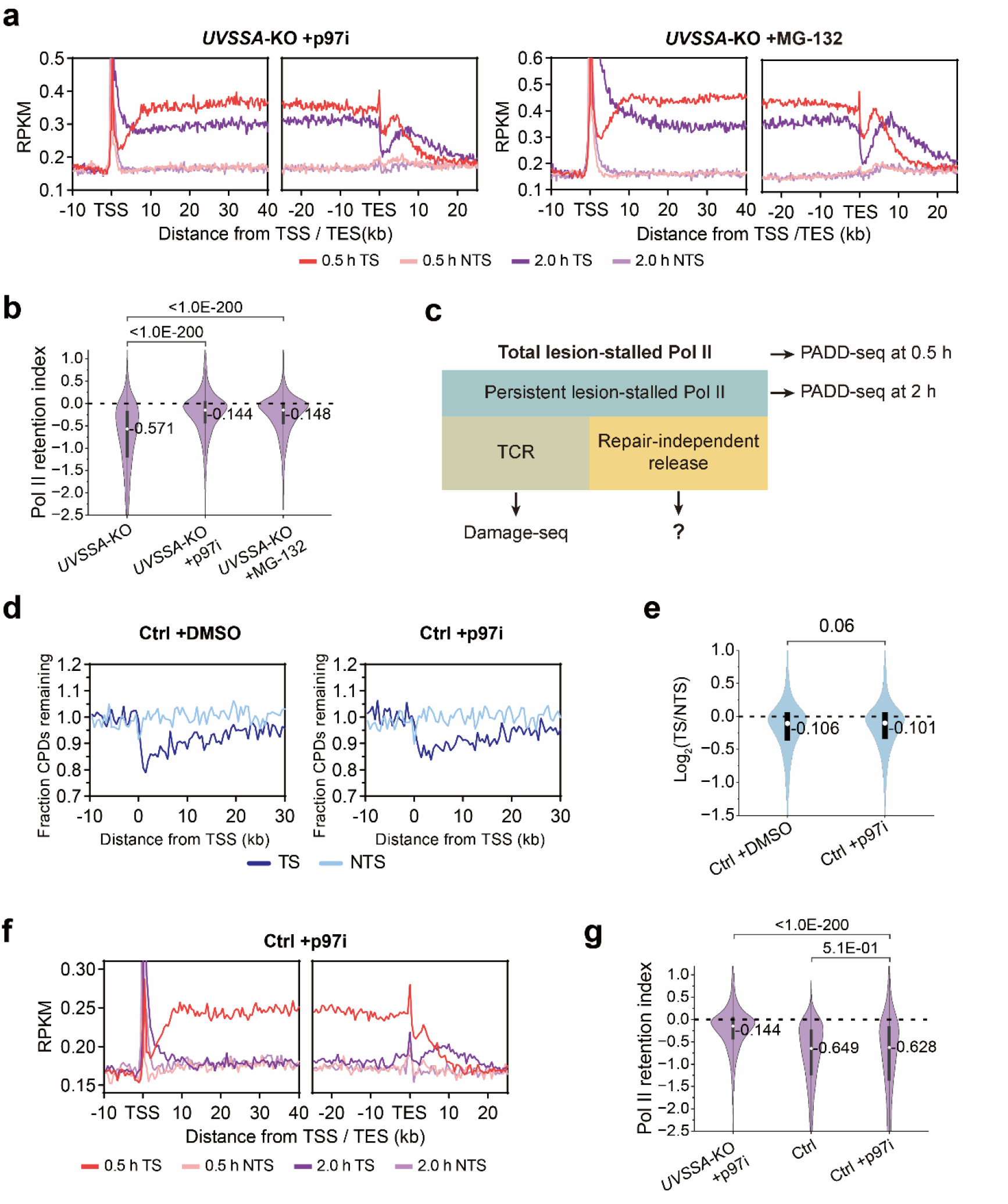
p97 extracts PolII from damage sites in the absence of UVSSA, while is dispensable for both TCR and PolII release in normal cells. **a,** Meta-gene analysis of PADD-seq signals around TSSs and TESs for active genes longer than 50 kb (*n* = 2790) under indicated conditions. **b,** Quantification of PolII retention on damage sites by relative change of PADD-seq signals from 0.5 h to 2 h on each gene. Active genes longer than 20 kb were selected (*n* = 4488). **c,** Diagram of three possible fates of lesion-stalled PolII in TCR-proficient cells. Total and persistent lesion-stalled Pol II are measured by PADD-seq at 0.5 h and 2 h, respectively, while TCR can be reflected by Damage- seq. **d,** Meta-gene analysis of Damage-seq signals around TSSs for active genes longer than 50 kb (*n* = 2790) under indicated conditions. Cells were collected immediately (0 h) or at 4 h after UV irradiation. Fraction CPDs remaining was calculated as the ratio of 4 h to 0 h. **e,** Violin plots of relative Damage-seq signals on each active gene (*n* = 6406). Log2 value of the ratio of fraction CPDs remaining on TS to that on NTS was calculated. **f,** As in **(a)** but for indicated conditions. **g,** As in **(b)** but for indicated conditions. *P* value was calculated using two-tailed paired Student’s *t*- test for **(b)**, (**e**) and **(g)**. See also Supplementary Fig. 3 and 4. Source data are provided as a Source Data file.

We next investigated whether p97 is also involved in TCR, and whether repair- independent PolII release driven by p97 occurs in TCR-proficient cells. When elongating PolII is blocked by a lesion, it can be resolved by TCR or removed by p97 independent of repair, or it can persist at the damage site (**Fig. 2c**). Since recruitment of TFIIH requires both UVSSA and PolII^17,44^, these three fates are incompatible with each other. We performed both Damage-seq and PADD-seq in XP-C cells in the presence of p97 inhibitor. Since prolonged p97 inhibitor treatment induces cell death (**Supplementary** Fig. 4a), CPD distributions at 0 h and 4 h post-UV were measured by Damage-seq and repair efficiency was determined by calculating the remaining fractions of damage on both strands. The results showed that inhibiting p97 had no significant impact on TCR (**Fig. 2d,e and Supplementary** Fig. 4b,c). Intriguingly, p97 inhibition also made no obvious difference on the disappearance of PolII-CPD interaction in TCR-proficient cells (**Fig. 2f,g and Supplementary** Fig. 4d-f). Thus, although inhibiting p97 should abolish repair-independent PolII release, the other two possible fates of stalled PolII, i.e., TCR and persistent stalling, were not significantly affected. The reasonable explanation is that the contribution of repair-independent PolII release in TCR-proficient cells is negligible. Furthermore, the fact that damage removal is not compromised by inhibiting p97 implies that p97 is not directly involved in TCR.

### UV-induced RPB1-K1268 ubiquitination plays important but not indispensable roles in both TCR and repair-independent PolII release

Multiple factors including PolII, CSB and UVSSA are ubiquitinated during TCR^14,15,27,45^. Among them, ubiquitination at K1268 of the PolII large subunit (RPB1- K1268ub) is of particular interest, since it was reported to be the major form of UV- induced PolII ubiquitination and involved in both TCR and UV-induced PolII degradation^15,27^. We generated RPB1-K1268R mutant (PKR) cells in which UV- induced PolII ubiquitination was almost abolished (**Supplementary** Fig. 5a,b), and assessed both TCR and PolII dissociation. Damage-seq revealed that the mutant cells had greatly reduced TCR compared to parental cells (**Fig. 3a,b**), consistent with a previous report^15^. However, in contrast to *CSB*-KO cells, preferential repair of transcribed strands in PKR cells was not completely eliminated (**Fig. 3a,b**). Since PKR cells are derived from GGR-deficient XP-C cells, this result suggests an important but not indispensable role of RPB1-K1268ub in TCR.

**Fig. 3.**
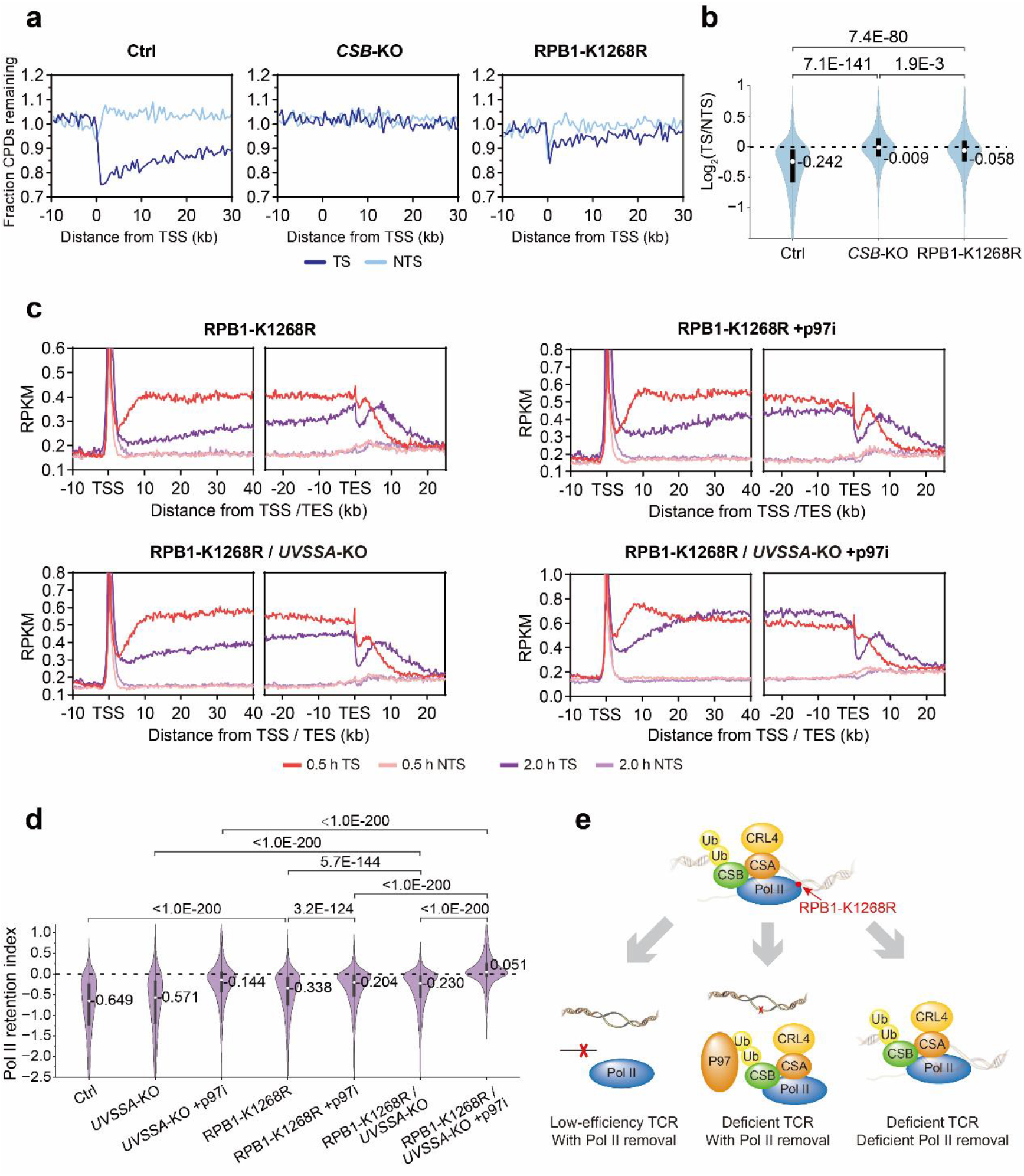
UV-induced RPB1-K1268 ubiquitination plays important but not indispensable roles in both TCR and repair-independent PolII release. **a,** Meta-gene analysis of Damage-seq signals around TSSs for active genes longer than 50 kb (*n* = 2790) under indicated conditions. Cells were collected immediately (0 h) or at 8 h after UV irradiation. Fraction CPDs remaining was calculated as the ratio of 8 h to 0 h. Data of WT and *CSB*-KO cells are from our previous study^35^. **b,** Violin plots of relative Damage-seq signals on each active gene (*n* = 6406). Log2 value of the ratio of fraction CPDs remaining on TS to that on NTS was calculated. **c,** Meta-gene analysis of PADD-seq signals around TSSs and TESs for active genes longer than 50 kb (*n* = 2790) under indicated conditions. **d,** Quantification of PolII retention on damage sites by relative change of PADD-seq signals from 0.5 h to 2 h on each gene. Active genes longer than 20 kb were selected (*n* = 4488). *P* value was calculated using two-tailed paired Student’s *t*-test for **(b)** and **(d)**. **e,** Diagram showing that three possible fates of lesion-stalled PolII should co-exist in RPB1-K1268R cells. See also Supplementary Fig. 5. Source data are provided as a Source Data file.

Unlike *CSA*-KO or *UVSSA*-KO cells, PolII retention was partially reduced at 2 h post UV in PKR cells (**Fig. 3c,d and Supplementary** Fig. 5c). Since lesion-stalled PolII could be resolved by either TCR or p97 in parental XP-C cells, we knocked out *UVSSA* (PKR/Uk double mutant cells, **Supplementary** Fig. 5d,e) or inhibited p97 in PKR cells to block each of these pathways, respectively. Under either condition, PolII retention at 2 h increased when compared to PKR cells without p97 inhibitor, however a portion of lesion-stalled PolII was still resolved (**Fig. 3c,d and Supplementary** Fig. 5c,f), suggesting that either TCR or p97-proteasome pathway could independently resolve lesion-stalled PolII with low efficiency in PKR cells. Blocking both pathways by inhibiting p97 in PKR/Uk cells abrogated the reduction of PolII-CPD interaction, further confirming that the partial resolution of lesion-stalled PolII in the absence of RPB1-K1268ub is still driven by TCR and p97-proteasome pathway (**Fig. 3c,d and Supplementary** Fig. 5f). These results suggest that although RPB1-K1268ub is an important target of p97 in repair-independent PolII release, p97 can also remove the TCR complex containing PolII from damage site with low efficiency in the absence of this modification. Thus, in contrast to parental XP-C cells in which TCR dominates, the three possible fates of lesion-stalled PolII, i.e., TCR, repair-independent PolII release and persistent lesion-stalled PolII, should co-exist in PKR cells (**Fig. 3e**).

### USP7 is involved in TCR but cannot abolish repair-independent PolII release driven by p97

The deubiquitinase USP7 is recruited to damage sites in complex with UVSSA during TCR, and is thought to play a role in TCR by deubiquitinating and stabilizing repair factors including CSB and UVSSA^12,46^. Consistent with previous reports, our Damage-seq data revealed that inhibiting USP7 significantly reduced TCR (**Fig. 4a,b and Supplementary** Fig. 6a,b). Under our experimental condition, the most affected repair factor is UVSSA and its mono-ubiquitination form (**Fig. 4c,d**), which was thought to play an important role in TCR^15^. Further inhibition of p97 could partially restore the levels of UVSSA and its mono-ubiquitination form in the chromatin fraction (**Fig. 4c,d**). However, the TCR activity was not significantly rescued by inhibiting p97 (**Fig. 4a,b and Supplementary** Fig. 6a,b), implying that USP7 promotes TCR not only by protecting repair factors from p97-proteasome degradation but also through other mechanisms.

**Fig. 4.**
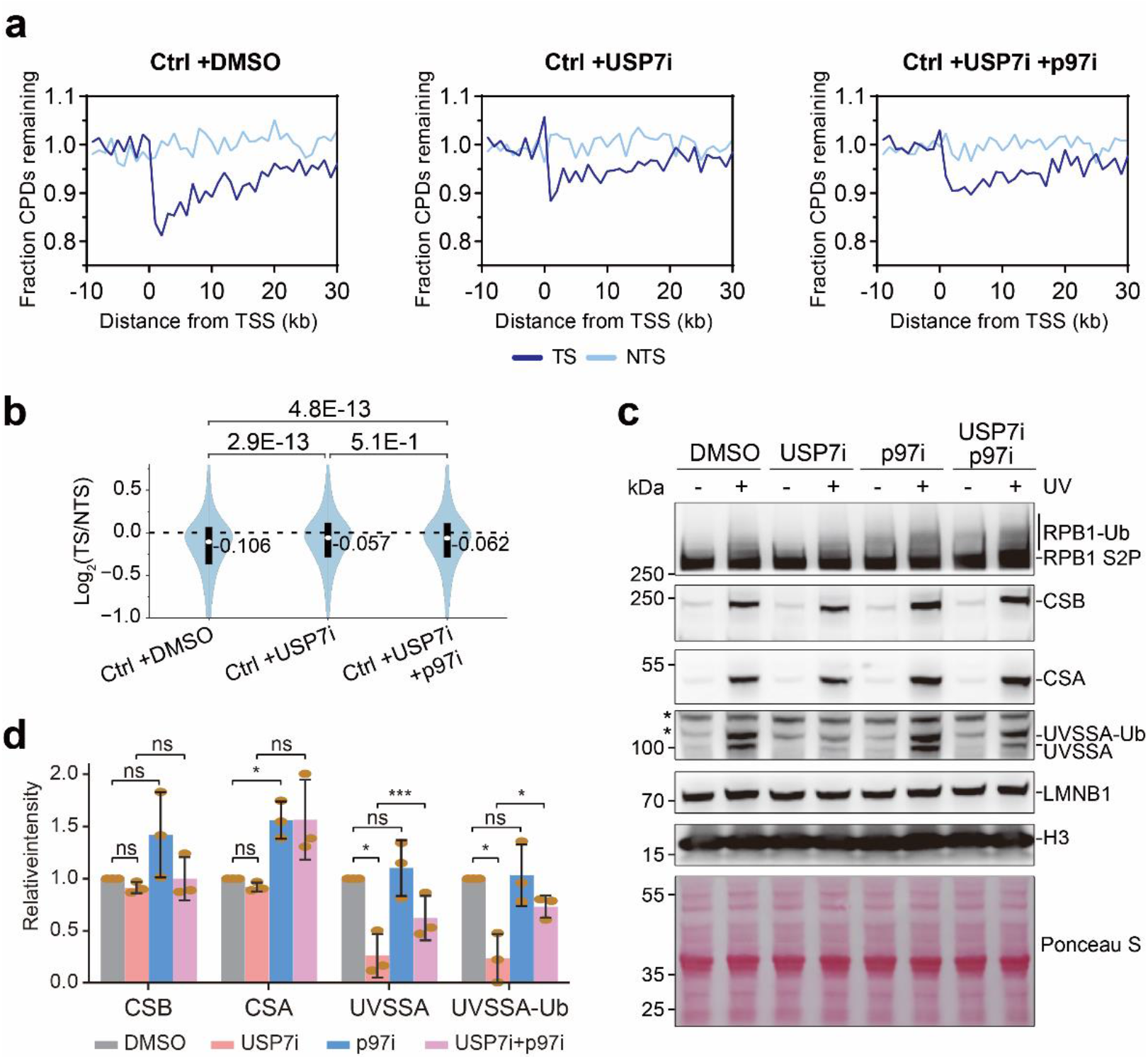
USP7 is involved in TCR. **a,** Meta-gene analysis of Damage-seq signals around TSSs for active genes longer than 50 kb (*n* = 2790) under indicated conditions. Cells were collected immediately (0 h) or at 4 h after UV irradiation. Fraction CPDs remaining was calculated as the ratio of 4 h to 0 h. **b,** Violin plots of relative Damage-seq signals on each active gene (*n* = 6406). Log2 value of the ratio of fraction CPDs remaining on TS to that on NTS was calculated. **c,** Western blot of chromatin fraction from XP-C cells treated with USP7i (FT671), p97i (CB-5083) or both at 2 h after 20 J/m^2^ UV-C treatment. The asterisks indicate two nonspecific bands that can also be seen in *UVSSA*-KO cells (Supplementary Fig. 1e). Experiments were performed in triplicate and a representative experiment is shown. **d,** Relative levels of CSB, CSA, UVSSA and UVSSA-Ub from three biological repeats. Data are shown as means with standard deviation. *P ≤ 0.05, **P ≤ 0.01, ***P ≤ 0.001. *P* value was calculated using two-tailed paired Student’s *t*-test for **(b)** and **(d)**. See also Supplementary Fig. 6. Source data are provided as a Source Data file.

Since the p97-proteasome pathway seems to have a negligible contribution to PolII removal in TCR-proficient cells (**Fig. 2f,g**), we investigated whether USP7, with its deubiquitinase activity, could prevent p97-mediated repair-independent PolII release. To disrupt TCR while maintaining the recruitment and deubiquitinase activity of UVSSA-USP7, we expressed the UVSSA-V411A mutant in *UVSSA*-KO cells, since the V411 residue is essential for its interaction with the p62 subunit of TFIIH but not required for UVSSA-USP7 interaction (**Fig. 5a,b**)^15,44,47^. Damage-seq showed that expression of UVSSA-WT, but not UVSSA-V411A, could efficiently rescue TCR (**Fig. 5c,d**). Accordingly, lesion-stalled PolII was completely resolved in the presence of UVSSA-WT, regardless of the presence of the p97 inhibitor (**Supplementary** Fig. 7a and 7b). By contrast, in UVSSA-V411A mutant cells the majority of PolII was removed from damage sites after 2 h with a minor portion of PolII remaining on lesions (**Fig. 5e top and 5f; Supplementary** Fig. 7c,d **top and 7e**). Treating with USP7 inhibitor resulted in complete clearance of lesion-stalled PolII in the mutant cells, indicating that USP7 could moderately reduce PolII release in TCR-deficient cells (**Fig. 5e middle and 5f; Supplementary** Fig. 7c **middle**). More importantly, inhibiting p97 could largely prevent PolII release in UVSSA-V411A mutant cells but not in cells expressing UVSSA-WT (**Fig. 5e bottom and 5f; Supplementary** Fig. 7a-d **bottom and 7e**), since the latter have efficient TCR. Therefore, while USP7 can partially impede p97-driven PolII release when TCR is absent, it cannot fully prevent this process.

**Fig. 5.**
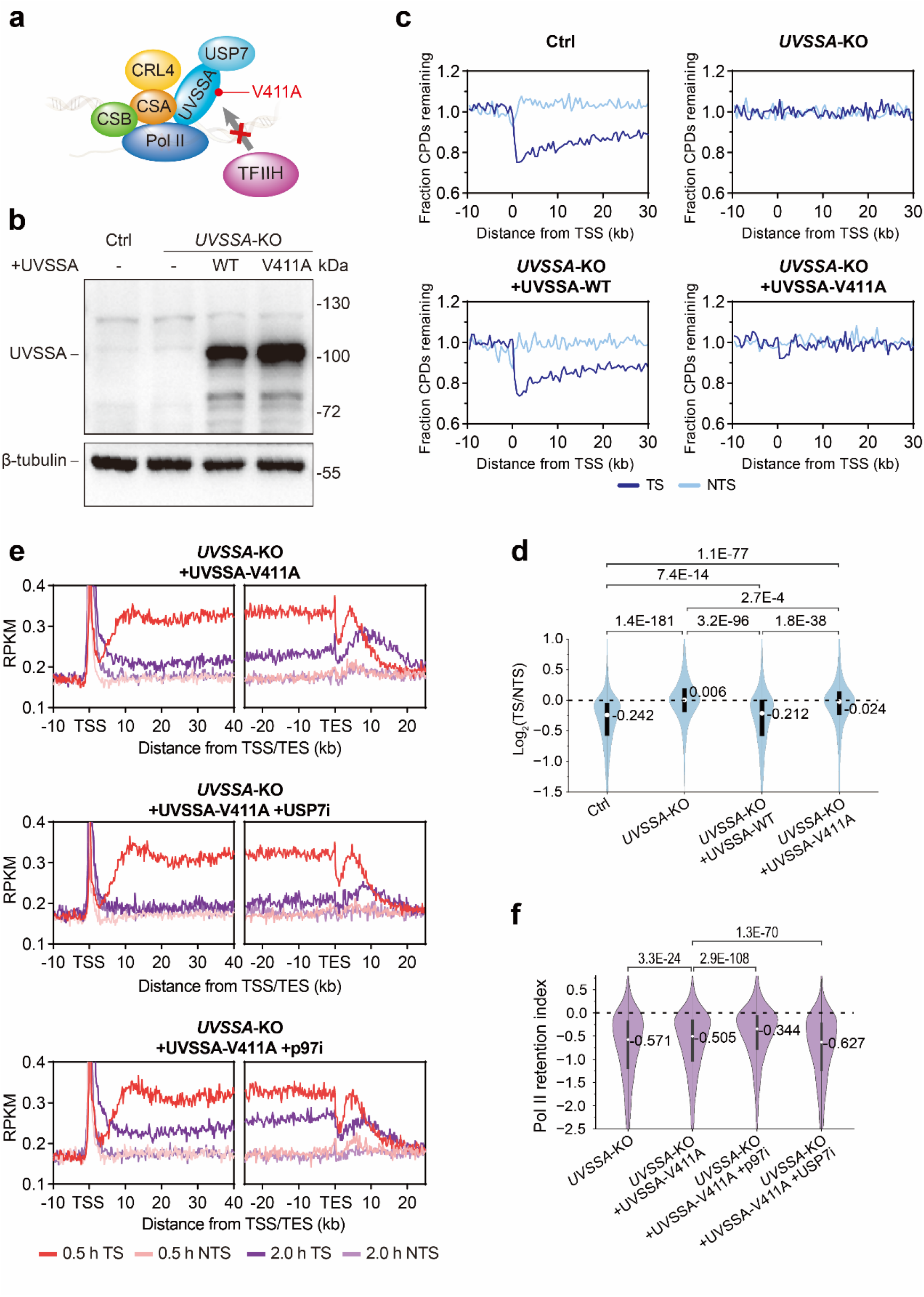
USP7 cannot abolish repair-independent PolII release driven by p97. **a,** A schematic showing that UVSSA-V411A mutation blocks the binding of UVSSA to TFIIH but not to USP7. **b,** Western blot of whole cell lysates showing the expression of UVSSA-WT or UVSSA-V411A in *UVSSA*-KO cells. Although the UVSSA antibody cannot detect endogenous UVSSA in whole cell lysates, it can detect ectopically expressed UVSSA. This experiment was performed once. **c,** Meta- gene analysis of Damage-seq signals around TSSs for active genes longer than 50 kb (*n* = 2790) under indicated conditions. Cells were collected immediately (0 h) or at 8 h after UV irradiation. Fraction CPDs remaining was calculated as the ratio of 8 h to 0 h. **d,** Violin plots of relative Damage-seq signals on each active gene (*n* = 6406). Log2 value of the ratio of fraction CPDs remaining on TS to that on NTS was calculated. **e,** Meta-gene analysis of PADD-seq signals around TSSs and TESs for active genes longer than 50 kb (*n* = 2790) under indicated conditions. **f,** Violin plots of relative change of PADD-seq signals from 0.5 h to 2 h on each gene. Active genes longer than 20 kb were selected (*n* = 4488). *P* value was calculated using two-tailed paired Student’s *t*-test for **(d)** and **(f)**. See also Supplementary Fig. 7. Source data are provided as a Source Data file.

## Discussion

Heritable defects of TCR-specific genes are related to clinically distinct genetic diseases. These include Cockayne syndrome (CS), a severe disorder characterized by compromised growth, impaired neural development and short life span, and UV sensitive syndrome (UV^S^S), a relatively mild disease with enhanced sensitivity to sunlight^5^. CS is caused by mutations in *CSA* or *CSB* genes^48^, while UV^S^S is mainly attributed to *UVSSA* mutations^11–13^ with a handful of exceptions that have mutations in *CSA*^49^ or *CSB*^50^. Persisting lesion-stalled PolII is thought to be more toxic than the damage itself and thus the cause of the severe symptoms of CS^15^. This hypothesis is supported by previous reports showing that PolII is restrained in damaged chromatin in UV-irradiated *CSA*- or *CSB*-defective cells but not in *UVSSA*-defective cells^11,51^. However, those conclusions were mainly drawn from Western blot or fluorescence recovery after photobleaching (FRAP) which could only detect total elongating PolII and its interaction with UV-irradiated chromatin, rather than the specific binding of PolII on UV-induced lesions^11,51^. Given that UV-induced lesions have a trans-effect of decreasing the elongation rate of PolII^32,33^, these technologies could not discriminate whether PolII retention in chromatin after damage was due to direct blocking or indirect slowing down of the polymerase. Therefore, we took advantage of PADD-seq to assess the direct PolII-damage interaction in TCR-deficient cells. We detected prolonged retention of PolII at damage sites in *CSB*- and *CSA*-KO cells, while this PolII-damage interaction disappeared within 2 hours in *UVSSA*-KO cells in a p97-dependent manner (**Figs. 1,2**), in agreement with two recent reports monitoring the association between PolII and damaged chromatin by FRAP or fluorescence microscope^52,53^. The phenomena are not only observed for UV-induced CPDs but also for cisplatin-adducts, although PolII are not completely removed from cisplatin-adducts in *UVSSA*-KO cells within 2 hours. The larger size and distinct blocking mechanism of cisplatin-adducts compared to UV-induced CPDs^34^ might differentially impact damage-induced PolII ubiquitination and subsequent eviction, resulting in incomplete removal of cisplatin-adducts-stalled PolII in *UVSSA*-KO cells.

Recent progress in structural biology revealed that TCR factors PolII-ELOF1- CSB-Cul4^CSA^-UVSSA form a complex with DNA damage^54^. Moreover, elongating PolII could interact with TFIIH after UV irradiation, indicating that PolII should not dissociate from damage sites before TFIIH loading^15,17^. Thus, TCR and repair-independent PolII release prior to UVSSA loading are mutually exclusive. This raised the question: how do cells coordinate these two competing processes to avoid the accumulation of lesion-stalled PolII while ensuring efficient TCR? The segregase p97 is a key factor in repair-independent PolII release, however, it is not required in TCR. Intriguingly, cullin-mediated ubiquitination is needed for TCR, implying that ubiquitination of PolII and other repair factors participates in TCR by regulating protein-protein interactions. TCR-proficient cells showed minimal p97-mediated removal of lesion-stalled PolII, however the role of p97 became prominent when TCR was partially or completely abrogated. These results suggested that the TCR pathway takes priority over the p97-driven repair-independent PolII release under normal conditions, while the p97-proteasome pathway works efficiently only when TCR is suppressed. A possible explanation is that the ubiquitin chains on PolII and other repair factors are involved in and masked by protein-protein interactions during efficient TCR and cannot be accessed by p97 (**Fig. 6**, **right**). Once the TCR process is interrupted, conformation change may happen to allow p97 to recognize and remove those ubiquitinated proteins (**Fig. 6**, **middle**). Only when both processes are blocked by deficient ubiquitination in *CSB*- or *CSA*-mutant cells does PolII persist on TBLs (**Fig. 6**, **left**).

**Fig. 6.**
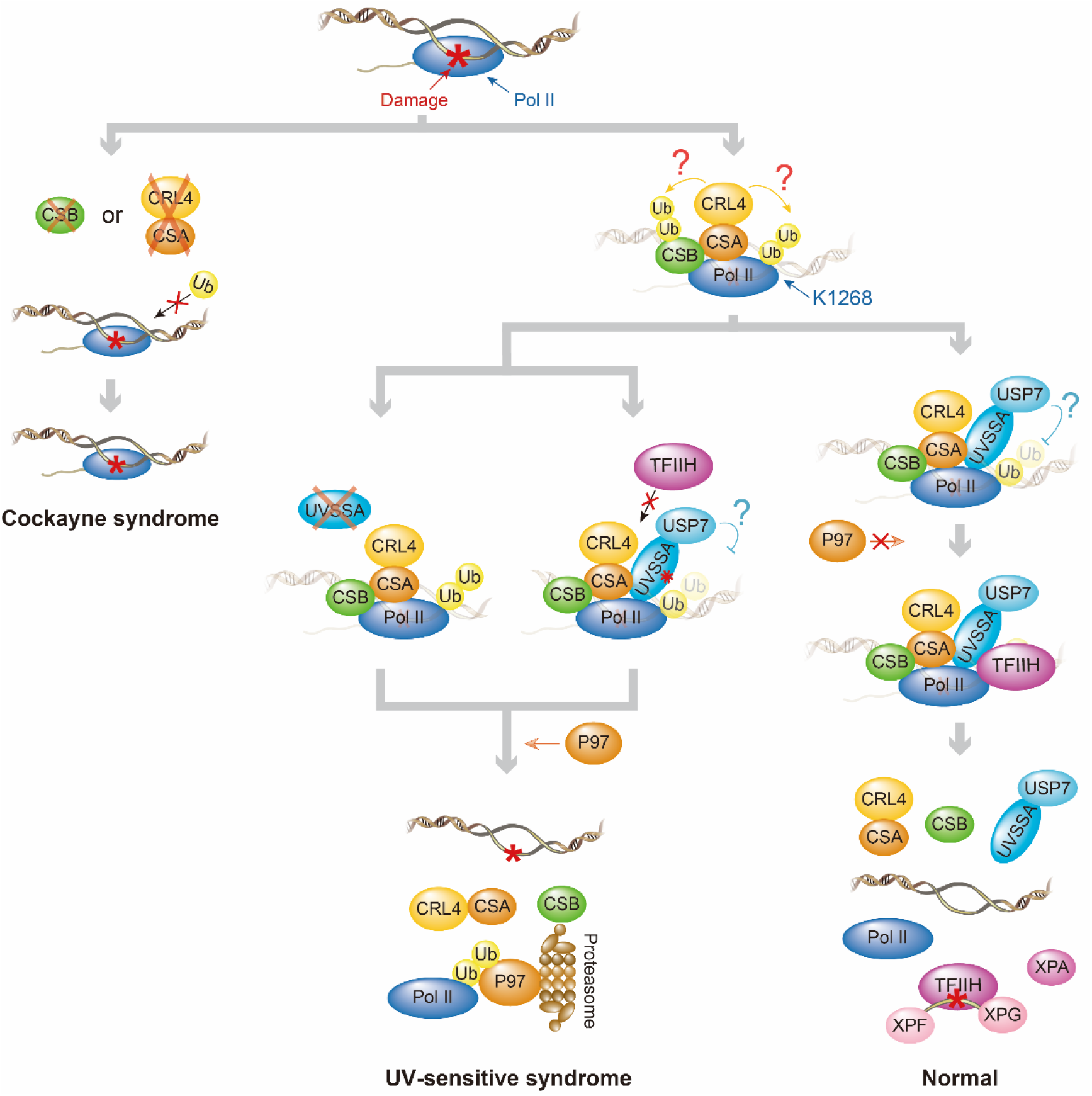
Working model of the coordination of TCR and repair-independent release of lesion-stalled PolII in response to TBLs. Left: For cells with deficiencies in *CSB* or *CSA* gene which are related to CS, PolII cannot be ubiquitinated, thus both TCR and p97-driven PolII release are blocked, resulting in extended retention of PolII on damage sites. Middle: For cells with mutated *UVSSA* which can cause UV^S^S, TCR is also blocked due to the lack of UVSSA or deficiency in TFIIH loading. However, PolII (and CSB) can be ubiquitinated, so p97 can remove lesion-stalled PolII to prevent its accumulation. Right: For normal cells with proficient TCR, p97 would not access ubiquitinated PolII during TCR, thus cannot interfere with repair. Lesion-stalled PolII should be resolved mainly by TCR.

Ubiquitination induced by TBLs is a complicated process. On one hand, the E3 ligase CRL4^CSA^ is a fundamental TCR factor^40^. On the other hand, the E3 ligases such as NEDD4^55^ and Cul5-based Elongin A complex^56^ are also reported to ubiquitinate PolII in this scenario. Nevertheless, our results show that either loss of CSA or the cullin family ubiquitin E3 ligases inhibitor could abolish both TCR and repair-independent PolII release. A recent study found that knocking down NEDD4 or Cul5 does not affect the dissociation of PolII from damaged chromatin^52^. Therefore, although there is UV-induced and CSA-independent PolII ubiquitination, CRL4^CSA^ is the most likely E3 ligase working on lesion-stalled PolII and required for both pathways. By contrast, although loss of UVSSA compromises UV-induced PolII ubiquitination, PolII is efficiently extracted from the damage site by p97, implying that lesion-stalled PolII can be ubiquitinated without UVSSA, albeit less efficiently. However, it is challenging to distinguish between ubiquitination on lesion-stalled PolII and ubiquitination on other PolII molecules.

Several important players of TCR including PolII, CSB, and UVSSA are ubiquitinated after UV irradiation^14,15,46^. Among them, ubiquitylation at K1268 of RPB1 subunit of PolII has been reported to play a key role in both TCR and UV-induced PolII pool regulation^15,27^. Although other lysine residues of RPB1 might also be ubiquitinated, only the K1268R mutation nearly abrogates UV-induced RPB1 ubiquitination while mutations of other lysine residues show no significant effect. Consistently, RPB1-K1268R mutation greatly reduces the efficiencies of both TCR and repair-independent PolII release under our conditions, but both processes were not completely blocked by this mutation. For TCR, loss of RPB1-K1268ub might compromise essential protein-protein interaction and diminish repair efficiency. In the case of repair-independent PolII release, RPB1-K1268ub is an important target of p97. However, in RPB1-K1268R mutant cells, other ubiquitinated repair factors, e.g. CSB, might also be recognized by p97, leading to the eviction of the whole complex, albeit with a reduced efficiency^14^. Nevertheless, the possibility that p97 recognizes the weak ubiquitination on other residues of RPB1 cannot be excluded. In agreement with this hypothesis, inhibition of cullin E3 ligase activity resulted in a stronger suppression of PolII release compared to RPB1-K1268R or RPB1-K1268R/UVSSA knockout. Thus, three scenarios described in **Fig.6** should co-exist in RPB1-K1268R mutant cells due to the low efficiency of either TCR or repair-independent release (**Fig. 3e**). Based on the elevated PolII stalling at damage sites, this mutation is expected to cause CS-like symptoms. However, no RPB1-K1268 mutation in patients has been reported to date. In a mouse model, this mutation is similar as *Csb*-KO that can cause CS in combination with *Xpa* deficiency, confirming our hypothesis. It is worth noting that in the absence of RPB1-K1268 ubiquitination, partial resolution of lesion-stalled PolII is not uniform along genes. Instead, the levels of persistent PolII gradually increased from TSSs to TESs, regardless of whether the resolution was driven by TCR or p97 (**Fig. 3c**). This trend coincides with phosphorylation of Ser2 (S2P) on RPB1-CTD that increases from TSSs to TESs along the gene body^39^. This increasing phosphorylation toward TESs might hinder the resolution of lesion-stalled PolII under certain conditions. However, there is currently no direct evidence, and further study is required to test this hypothesis.

Remarkably, there is not only a ubiquitin E3 ligase (CRL4^CSA^) but also a deubiquitinase complex, namely UVSSA-USP7, among the fundamental TCR factors. It was assumed that USP7 could protect repair factors from proteasome degradation and promote TCR with its deubiquitinase activity^12,46^, whereas UVSSA plays a more important and direct role in TCR by recruiting TFIIH through its interaction with p62 subunit^15,17,44^. Our results showed that inhibiting the deubiquitinase activity moderately suppressed TCR, however, inhibition of p97 could not significantly rescue repair, indicating that maintaining the stability of repair factors is not the sole mechanism by which USP7 promotes TCR. It is possible that both ubiquitination and deubiquitylation can regulate protein-protein interactions to promote TCR, but the effect of USP7 inhibition is weaker than cullin inhibition. On the other hand, UVSSA-V411A mutation slightly impeded p97-driven PolII release and this impediment was eliminated by inhibiting USP7, suggesting that the deubiquitinase activity of USP7 just marginally prevents PolII release when TCR is blocked. Therefore, although no actual case has been reported, it is expected that the UVSSA-V411A mutation should not cause severe CS-like symptom as RPB1-K1268R mutation. Thus, p97 may play a broader role in preventing the accumulation of lesion-stalled PolII when TCR is blocked in a late stage after the recruitment of UVSSA-USP7.

Impaired clearance of TBLs induced by endogenous factors like formaldehyde is thought to be the underlying cause for the severe symptoms of TCR deficiency in CS^57^. However, TCR deficiency might not be the direct cause of CS, since only mutations in *CSA* and *CSB* genes lead to CS^48^, while mutations in *UVSSA* result in significantly milder symptoms, although they all completely abrogate TCR^11–13^. Here we uncovered different fates of lesion-stalled PolII in cells lacking CSA/CSB compared to UVSSA, implying an association between persistent lesion-stalled PolII and the severe symptoms of CS. If TCR is defective while lesion-stalled PolII can be released, lesions can be removed by GGR or other pathways^58^, or bypassed during replication. The cells are still sensitive to UV, but affected individuals exhibit only symptoms of UV^S^S. In contrast, if TBLs are tightly bound by stalled PolII, they could not be handled by any other mechanism, and the patients would develop CS (**Fig. 6**). In addition, the phenotype of TCR-deficient mutations might also be affected by other pathways and genes. It was reported that similar *CSB*-null mutations can cause different phenotypes including UV^S^S, mild and severe CS^50,59,60^. It is possible that other genes involving in various pathways such as alleviating PolII retention, reducing endogenous DNA damage formation, and compensating for the detrimental consequences of persistent lesion-stalled PolII, might also contribute to the symptoms. However, the exact underlying mechanisms for specific patients are unclear. Moreover, although deficiencies in both GGR-specific genes and common NER genes can result in xeroderma pigmentosum (XP) with high risk of skin cancer, only mutations in common NER genes that also impair TCR can cause neurodegeneration or even CS-like symptoms in addition to XP, albeit usually milder than typical CS^24^, implying a similar mechanism underlying these symptoms and CS phenotype induced by mutations of TCR-specific genes. Our results suggested the possibility that UVSSA-USP7 could hinder the clearance of lesion-stalled PolII in some NER-deficient cells, and the remaining lesion-stalled PolII might cause relatively mild neurodegeneration or CS-like symptoms. Recent studies reported that some patients with onset neurodegeneration symptoms had mutations in common NER genes including *XPA*, *XPD* and *XPF*, but they only showed elevated sensitivity to sunlight rather than typical XP symptoms^61^. Furthermore, a recent paper demonstrated that physiological natural aging is related to increased PolII stalling^62^, which is also observed in pathological premature aging caused by CS. Endogenous DNA damage is thought to be the reason for increased PolII stalling during aging. Therefore, persistent lesion-stalled PolII might be a common cause in neurodegeneration and aging rather than just the underlying reason of CS, and targeting lesion-stalled PolII for degradation might be a potential way to relieve neurodegeneration in patients with TCR deficiency. In summary, our study unveiled the mechanism to regulate the fate of lesion-stalled PolII, shedding light on potential therapeutic strategy of neurological symptoms caused by transcription-blocking damage.

## Methods

### Cell culture

Human skin fibroblast cells derived from an XP-C patient (GM15983, dubbed “XP-C cells”) were purchased from Coriell Institute. Cells were grown in DMEM (Gibco) supplemented with 10% fetal bovine serum (Gibco) and 1% Penicillin–Streptomycin (Gibco) at 37 °C in a 5% CO2 humidified chamber.

### Gene editing by CRISPR-Cas9

*CSA* and *UVSSA* knock-out cells and RPB1-K1268R mutation cells were generated by the CRISPR- Cas9 mediated gene editing strategy^63^. For the gene deletion experiment, the sgRNA coding sequence **(Supplementary Table 1)** was cloned into pX459 V2.0 vector (a gift from Feng Zhang; Addgene plasmid 62988)^63^. The plasmid was transfected into designated cells using HighGene transfection reagent (ABclonal). 24 h after transfection, cells were selected for 24 h with 1 μg/ml puromycin (Selleck). Single clones were isolated from surviving fraction by limiting dilution. The sgRNA target regions were amplified by PCR and the products were ligated to the pEASY-Blunt Zero vector (TransGen Biotech). The PCR products and recombinant vectors were sequenced by Sanger sequencing. Knock-outs were further confirmed by Western blot.

Generation of RPB1-K1268R mutants were achieved by inducing site-specific double-strand breaks near the designated RPB1 lysine residues in combination with homology-directed-repair oligonucleotides carrying the K1268R substitutions. A ∼100 nt oligonucleotide **(Supplementary Table 1)** was co-transfected with a sgRNA expressing plasmid corresponding to the target sequence. The introduction of designated amino acid substitution was confirmed by Sanger sequencing and loss of UV-induced RPB1 ubiquitination (see section of *Detection of ubiquitylated RPB1*).

### Retrovirus production and generation of UVSSA-V411A cell line

The wild-type human UVSSA cDNA was fused with a N-terminal FLAG-HA tag. UVSSA-V411A mutant with the N-terminal FLAG-HA tag was generated by site-directed PCR mutagenesis using specific primer sets (**Supplementary Table 2**), 2×Phanta Max Master Mix (Vazyme) and the DpnI restriction enzyme (Sangon Biotech). Then the wild-type and V411A mutational UVSSA constructs were cloned into pMXs-IRES-Puro retroviral expression vector (a gift from Dr. Feilong Meng, Chinese Academy of Sciences, Shanghai, China), respectively. For retrovirus production, 293T cells were transfected with the UVSSA-encoding plasmids constructed above together with packaging plasmid PCL10A1 (a gift from Dr. Feilong Meng) using HighGene transfection reagent (Abclonal). Viral particles were collected 48 h after transfection, filtered through 0.45-μm filters, and infected into *UVSSA*-KO XP-C cells. After incubation for 48 h, cells were selected with 1 μg/ml puromycin for 24 hours. Surviving fractions were further incubated for 48 h in DMEM supplemented with 20% conditioned medium, 15% fetal bovine serum and 1% penicillin-streptomycin. All the constructs and mutants were confirmed by Sanger sequencing.

### UV irradiation and drug treatments

Cells were prepared in Petri dishes and treated with 2 μM MLN-4924 (MedChemExpress) as CRL inhibitor (CRLi), 10 μM FT671 (MedChemExpress) as USP7 inhibitor (USP7i), 5 μM CB-5083 (MedChemExpress) as p97 inhibitor (p97i), 50 μM MG-132 (Selleck) as proteosome inhibitor, or 250 nM NVP-2 (MedChemExpress) as CDK9 inhibitor, as indicated. After incubation at room temperature for 5 min, the media was removed and cells were irradiated with 20 J/m^2^ 254 nm UV-C. Then the drug-containing culture medium was added back, and cells were incubated for the indicated time courses at 37 °C in a 5% CO2 humidified chamber.

### Detection of TCR factors in chromatin fractions

XP-C cells were prepared in 60-mm plates. UV irradiation and drug treatment of USP7i and p97i were performed as described above, followed by the incubation for 2 h at 37°C in a 5% CO2 humidified chamber. Then cells were harvest by trypsinization and centrifugation.

Cells were lysed in 200 μl of lysis buffer 1 (50 mM HEPES-KOH pH 7.5, 1 mM EDTA, 140 mM NaCl, 0.25% Triton X-100, 0.5% CA-630 and 10% glycerol) supplemented with protease/phosphatase inhibitor cocktail (Roche) for 20 min on ice. The pellets were collected by centrifugation at 20,000 g at 4°C for 3 min, and sequentially washed with 100 μl of lysis buffer 1 and 100 μl of lysis buffer 2 (10 mM Tris-HCl pH 8.0, 1 mM EDTA, 200 mM NaCl and 0.5 mM EGTA) supplemented with protease/phosphatase inhibitor cocktail (Roche) very gently. Then the pellets were incubated with 5 μl of Super Nuclease (Smart-Lifesciences) in 50 μl of Buffer 3 (50 mM HEPES-KOH pH 7.5, 140 mM NaCl, 0.25% Triton X-100, 0.5% CA-630 and 10% glycerol) supplemented with protease/phosphatase inhibitor cocktail (Roche) for 15 min on ice, and boiled at 96-98°C for 10 min with the supplement of SDS-PAGE Protein Sample Loading Buffer (Beyotime). Samples were span down and the supernatants were resolved by 4-15% SDS-PAGE. The CTD-Ser2- phosphorylated RPB1, CSB, CSA, UVSSA and H3 (loading control) were detected by indicated antibodies (see *Western blot* section for details). Experiments were performed in triplicate. Protein levels were quantified and normalized to signals of Ponceau S staining. *P* value was calculated using two-tailed paired Student’s *t*-test.

### Detection of ubiquitylated RPB1

Dsk2-pulldown of ubiquitylated RPB1 were performed as described previously^64^. To prepare Dsk2- coated beads, purified GST-Dsk2 protein (a gift from Dr. Fenglong Meng) was incubated with pre-washed glutathione agarose beads [2 ml of original beads suspension (Smart-Lifesciences) was washed with PBSA (1xPBS plus 1% BSA) supplemented with 2 mM DTT] at a rotator overnight at 4°C. The beads were span down, washed twice with cold PBSA containing protease inhibitor and 0.1% Triton X-100, and then washed once with PBSA containing protease inhibitor. The prepared Dsk2-coated beads were resuspended in 20 ml of PBSA containing protease inhibitor and 0.02% sodium azide and stored at 4°C.

WT and RPB1-K1268R cells cultured in 60-mm dishes were irradiated with or without 20 J/m^2^ UV-C, and incubated for 30 min at 37 °C in a 5% CO2 humidified chamber. After the incubation, cells were harvested by trypsinization and centrifugation, and lysed in 200 μl of TENT buffer (50mM Tris-HCl pH7.4, 2mM EDTA, 150mM NaCl, 1% Triton X-100) containing inhibitors of protease and phosphatase for 10 min on ice. Then the samples were sonicated by the sonicator (Qsonica) at 30% amplitude for 7 min with 30 s ON and 30 s OFF pulses at 4°C. Samples were centrifuged at 20,000 g for 5 min at 4°C to remove the debris, and the supernatants were saved. Protein concentration was quantified by the A280 (absorbance at 280 nm) using a spectrophotometer (DeNovix). Equal amounts of supernatants were saved as Input.

Dsk2-coated beads were pre-washed twice in TENT buffer. Each 0.4 ml of Dsk2-coated beads suspension (equivalent to 10 μl of packed beads) was used to pull down ubiquitylated RPB1 from equal amounts (less than 1 mg) of whole cell extracts. Samples were incubated in 200 μl of TENT buffer containing protease inhibitor and phosphatase inhibitor on a rotator at 4°C overnight. The beads were washed three times with TENT buffer and centrifuged at 750 g for 1 min at 4°C, and then all liquid was removed. The beads and Input samples were then boiled at 96-98°C for 10 min in SDS-PAGE Protein Sample Loading Buffer. Samples were span down and the supernatants were resolved by 4-20% SDS-PAGE. The CTD-Ser2-phosphorylated RPB1, ubiquitin and β-tubulin (loading control) were detected by indicated antibodies (see *Western blot* section for details).

### Western blot

Samples were prepared as described above and resolved by precast 4-15% or 4–20% gradient gels for SDS-PAGE (Beyotime). Resolved protein samples were transferred to nitrocellulose membranes (PALL), followed by Ponceau S (Sigma) staining and blocking for 1 h at room temperature in 5% skim milk in TBST (50 mM Tris-HCl pH 7.6, 150 mM NaCl, 0.1% Tween 20). The membranes were incubated with indicated primary antibodies in 5% BSA in TBST overnight at 4°C. Membranes were washed three times in TBST, followed by incubation with 1:5000 diluted HRP-conjugated secondary antibodies (Beyotime: anti-mouse, A0216; anti-rabbit, A0208) in 5% skim milk in TBST for 1 h at room temperature. After extensive washing with TBST, the proteins were visualized using enhance chemiluminescence reagent (Tanon). Primary antibodies used for immunoblotting: mouse anti-β-tubulin monoclonal antibody, 1:1000 (Cell Signaling Technology, 86298); rabbit anti-CTD- Ser2-phosphorylated RPB1 polyclonal antibody, 1:1000 (Abcam, ab5095); rabbit anti-CSA monoclonal antibody, 1:1000 (Abcam, ab137033); rabbit anti-CSB polyclonal antibody, 1:1000 (Bethyl Laboratories, A301-345A); mouse anti-UVSSA polyclonal antibody, 1:500 (Abnova, H00057654-B01P); rabbit anti-H3 monoclonal antibody, 1:1000 (Cell Signaling Technology, 4499); mouse anti-ubiquitin monoclonal antibody, 1:200(santa cruz, sc-8017); mouse anti-Lamin B1 monoclonal antibody, 1:1000 (santa cruz, sc-377000).

### Cell viability

XP-C cells were equally seeded in 6-cm culture dishes, and the cell density was controlled at about 50%. The second day, cells were treated with 20 J/m^2^ UV-C and 5 μM CB-5083 (or vehicle DMSO), and incubated for 0 h, 4 h, or 8 h. Cells were carefully washed with PBS, fixed with 100% methanol for 10 min, stained with 0.5% (w/v) crystal violet (Sangon Biotech) in 25% methanol for 10 min at room temperature, and washed with ddH2O. Pictures were taken by a normal camera (left) or a microscope (right).

### Damage-seq

Cells cultured in 60-mm plates were subjected to UV irradiation and drug treatment of CRLi, USP7i and p97i as described above, followed by incubation for 0 h (no incubation), 4 h or 8 h at 37°C in a 5% CO2 humidified chamber. Then cells were scraped and collected by centrifugation. Genomic DNA was extracted using PureLink Genomic DNA Mini Kit (Thermo Fisher Scientific), and sonicated by a Q800 Sonicator to get DNA fragments averagely 300 - 600 bp in length. The 500 ng of DNA fragments were used for Damage-seq^37^. Briefly, DNA fragments were subject to end-repair and dA-tailing with NEBNext Ultra II DNA Library Prep Kit for Illumina (New England Biolabs), followed by ligation with 50 pmol of adaptor 1 (Ad1, **Supplementary Table 3**) at 4 °C overnight. Samples were purified by 0.8×DNA FragSelect XP Magnetic Beads (Smart-lifesciences) and used for immunoprecipitation with the anti-CPD antibody (Cosmo Bio, NMDND001). Then, primer O3P (**Supplementary Table 3**) was attached to IP-purified DNA and was extended by NEBNext Ultra II Q5 Master Mix (New England Biolabs), followed by ExoI (New England Biolabs) treatment and 1.1×DNA FragSelect XP Magnetic Beads cleanup. Purified extension products were denatured and ligated to adaptor 2 (Ad2, **Supplementary Table 3**) by Instant Sticky-end Ligase Master Mix (New England Biolabs) at 4 °C overnight. Ligation products were purified by 1.1×DNA FragSelect XP Magnetic Beads and amplified by 12-15 cycles of PCR with NEBNext Ultra II Q5 Master Mix and NEBNext Multiplex Oligos for Illumina (New England Biolabs) according to manufacturer’s instructions. Libraries were sequenced from both ends on an Illumina NovaSeq system by Mingma Technologies Company to get ∼40 million raw reads for each sample.

### PADD-seq

PADD-seq was performed as described previously^35^. Briefly, cells cultured in three 150-mm plates were used for each sample. For PADD-seq experiments of PolII and CPD, UV irradiation and drug treatment of NVP-2, CRLi, p97i, USP7i and MG-132 were performed as described above, followed by the incubation for 0.5 h or 2 h at 37°C in a 5% CO2 humidified chamber. For PADD- seq experiments of PolII and cisplatin-adduct: cisplatin (Sigma) was dissolved in DMSO to 20 mM and immediately added to medium to a final concentration of 200 μM. Cells were treated with 5 μM CB-5083 (p97i) together with cisplatin in some experiments, as indicated in the figure legends. Then cells were incubated at 37 °C for 1.5 h, washed twice with PBS, and treated with 250 nM NVP-2 (or together with 5 μM CB-5083), followed by further incubation for 0.5 h or 2 h at 37°C in a 5% CO2 humidified chamber.

Then the cross-linking was performed by incubation with 1% formaldehyde (Thermo Fisher Scientific) for 10 min at room temperature with gentle shaking, and stopped by incubation with 150 mM Glycine (Sigma) for 5 min at room temperature. Cells were washed twice by ice-cold PBS, scraped and collected by centrifugation. Collected cells were lysed in lysis buffer 1 (50 mM HEPES-KOH pH 7.5, 1 mM EDTA, 140 mM NaCl, 0.25% Triton X-100, 0.5% CA-630 and 10% glycerol) supplemented with protease inhibitor cocktail (Roche) on ice for 10 min. Pellet was collected by centrifugation, followed by incubation with lysis buffer 2 (10 mM Tris-HCl pH 8.0, 1 mM EDTA, 200 mM NaCl and 0.5 mM EGTA) supplemented with protease inhibitor cocktail on ice for 10 min. The pellet was collected by centrifugation and resuspended in lysis buffer 3 (10 mM Tris-HCl pH 8.0, 1 mM EDTA, 140 mM NaCl, 1% Triton X-100, 1.5% SDS and 0.1% Na-DOC) supplemented with protease inhibitor cocktail on ice for 30 min. Obtained chromatin lysate was sonicated by a Q800 Sonicator (Qsonica) to get DNA fragments averagely 300 - 600 bp in length, followed by centrifugation at 20,000g for 10 min at 4 °C to collect the supernatant. Sample concentrations were determined by Qubit dsDNA BR Assay kit (ThermoFisher Scientific).

Fragmented chromatin was subjected to chromatin immunoprecipitation. In brief, 167 μg of fragmented chromatin and 25 μg of anti-RPB1 antibody (Bethyl Laboratories, A304-405A) were incubated in RIPA buffer (10 mM Tris-HCl pH 8.0, 1 mM EDTA, 140 mM NaCl, 1% Triton X-100, 0.1% SDS and 0.1% Na-DOC) supplemented with protease inhibitors, 0.1 % BSA (Sigma) and 0.1 μg/μl tRNA (Sigma) for 2 h on a rotator at 4 °C. Then 60 μl of protein A agarose beads (Smart-lifesciences) were added and the mixture was incubated on a rotator at 4 °C overnight. Beads were sequentially washed with RIPA buffer, RIPA-500 buffer (10 mM Tris-HCl pH 8.0, 1 mM EDTA, 500 mM NaCl, 1% Triton X-100, 0.1% SDS and 0.1% Na-DOC), LiCl Wash buffer (10 mM Tris-HCl pH 8.0, 1 mM EDTA, 250 mM LiCl, 0.5% CA-630 and 0.5% Na-DOC) three times for each buffer and once with TE (10 mM Tris-Cl pH 8.0 and 1 mM EDTA), followed by elution with Direct Elution buffer (10 mM Tris-HCl pH 8.8, 5 mM EDTA, 300 mM NaCl and 1% SDS). Elutes were treated with RNase A (ThermoFisher Scientific) for 30 min at 37 °C, followed by incubation with proteinase K (Takara) for 2 h at 55 °C. Then cross-linking was reversed by incubation at 65 °C overnight. DNA was purified by phenol-chloroform extraction and ethanol precipitation, and the concentration was determined by Qubit dsDNA HS Assay Kits (ThermoFisher Scientific).

Purified DNA (50-100 ng) was subjected to Damage-seq using an anti-CPD antibody (for PADD-seq of PolII and CPD) or an anti-cisplatin-adduct antibody (Abcam, ab103261, for PADD- seq of PolII and cisplatin-adduct) as previously described^35,37,65^. Libraries were sequenced from both ends on an Illumina NovaSeq system by Mingma Technologies Company to get ∼20 million raw reads for each sample.

### Genome alignment and visualization

For Damage-seq and PADD-seq, reads containing the Ad1 sequence at the 5’ end were removed using Cutadapt (version 1.12). Reads were further trimmed by trim_galore (version 0.6.7) and then aligned to the reference genome hg38 using BWA MEM (version 0.7.17)^66^ with default parameters. Sambamba (version 0.8.1)^67^ and in-house Python scripts were applied to remove unmapped reads, duplicate reads, unpaired reads, reads with a mapping quality of <25 and reads with a secondary alignment. The damage sites (CPD or cisplatin-adduct) for Damage-seq and PADD-seq are expected to be the two nucleotides upstream of mapped reads. Reads with relative dinucleotide (TT, TC, CT and CC for CPD; GG and AG for cisplatin-adduct) at expected damage sites were selected.

Active genes [fragments per kilobase of exon per million reads mapped (FPKM)>1] were selected using the gene quantification data of human foreskin fibroblast cells (BJ) obtained from the ENCODE consortium (accession number ENCSR000COP). Genome annotation was obtained from Ensembl at http://www.ensembl.org/, and active genes >2 kb apart from each other were selected using Bedtools (version 2.30.0)^68^.

Strand-specific bedgraph files were generated using the bamCoverage tool of deepTools2^69^ and normalized to reads per kilobase per million reads mapped (RPKM) values. Screenshots were plotted using IGV (version 2.9.2)^70^. Meta-gene analyses were performed using in-house Python scripts.

For quantification of PADD-seq signals on each gene: Active genes longer than 20 kb were selected (*n* = 4488). Signals from TSS downstream 10 kb to TES were calculated for each gene using multiBamSummary tool of deepTools2 and normalized to RPKM values. To describe the change of PADD-seq signals on each gene from 0.5 h to 2 h timepoint, we defined the Pol II retention index as follows: Pol II retention index = [(TS2 h-NTS2 h)-(TS0.5 h-NTS0.5 h)]/[AVERAGE(TS0.5 h- NTS0.5 h)]. The negative value of Pol II retention index represents that PolIIs were released from lesions, while the 0 or positive value represents PolIIs were restrained at lesions.

For quantification of Damage-seq signals on each gene: Signals on each active gene (*n* = 6406) were calculated using multiBamSummary tool of deepTools2. Because GGR is deficient in XP-C cells, the repair of total CPDs is negligible. Thus, fraction CPDs remaining can be calculated as the ratio of 4 h or 8 h to 0 h. Log2 value of the ratio of fraction CPDs remaining on TS to that on NTS was calculated.

## Statistics and reproducibility

Statistical analyses were performed with a two-tailed paired *t*-test on Microsoft Excel. The exact value of *n*, representing the number of genes, is indicated in the figure legends. The assays were performed once or twice as indicated. The investigators were not blinded to allocation during experiments and outcome assessment, as all analyses were objective in nature.

## Reporting summary

Further information on research design is available in the Nature Portfolio Reporting Summary linked to this article.

## Data availability

Raw sequencing data for PADD-seq and Damage-seq is publicly available at NCBI Sequence Read Archive under accession number PRJNA1074553. Source data are provided with this paper. Further information, resources, and reagents are available from the corresponding author upon request.

## Code availability

The codes are publicly available at https://github.com/Huulab/Analysis-of-Damage-seq-and-PADD-seq.

## Acknowledgements

This work was supported by National Key R&D Program of China [2022YFA1303000], National Natural Science Foundation of China (NSFC) [32271343], Shanghai Municipal Natural Science Foundation [22ZR1413900], innovative research team of high-level local university in Shanghai (to J.H.) and Israel Science Foundation grant 482/22 (to S.A.). And S.A. is the recipient of the Jacob and Lena Joels memorial fund senior lectureship. The funders had no role in study design, data collection and analysis, decision to publish or preparation of the paper. This work was supported by the Medical Research Data Center of Fudan University. We thank Dr. Feilong Meng (Center for Excellence in Molecular Cell Science, Chinese Academy of Sciences, Shanghai, China) for providing the pMXs-IRES-Puro retroviral expression vector, packaging plasmid PCL10A1, and GST-Dsk2 protein.

## Author contributions

J.H., S.A. and Y.Z. conceived and coordinated the project. X.Z., Y.H., Y.T., Y.Z. and S.W. generated cell lines. Y.Z., X.Z., M.G., Y.H. and D.Z. performed PADD-seq and Damage-seq. M.G., X.Z. and Y.T. carried out all other experiments. J.H., S.A., Y.Z. and A.P. analyzed data. J.H., Y.Z., X.Z. and M.G. wrote the original draft of the paper. J.H., Y.Z., S.A. and A.P. revised the paper. J.H. supervised the study.

