## Supplementary Information for "Coordination of transcription-coupled repair and repair-independent release of stalled RNA polymerase II in response to transcription-blocking lesions"

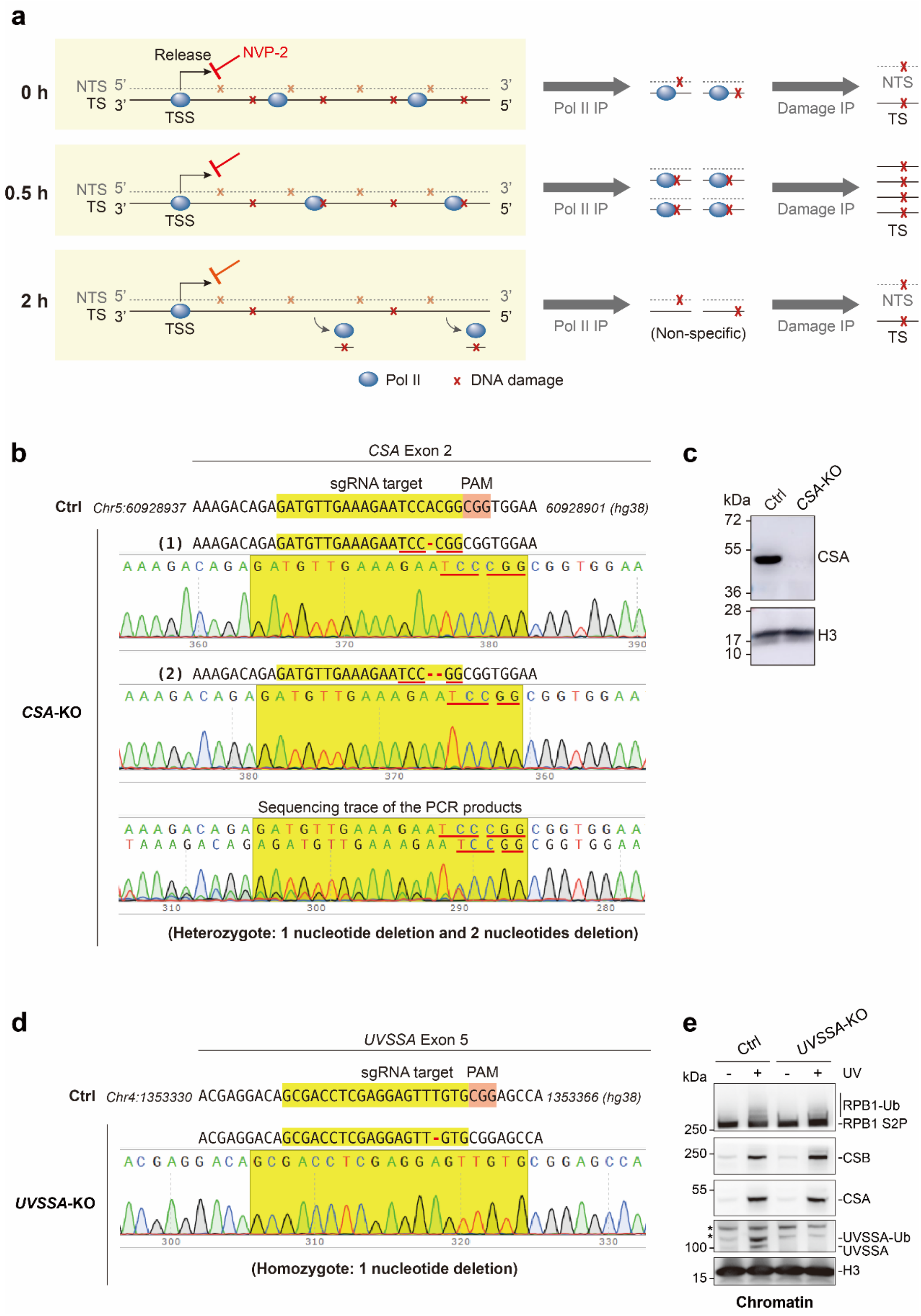
a
0 h
0.5 h
2 h
Release
NVP-2
NTS 5'
TS 3'
3'
5'
TSS
Pol II IP
Damage IP
NTS
TS
(Non-specific)
Pol II
DNA damage
b
CSA Exon 2
sgRNA target
PAM
Ctrl Chr5:60928937 AAAGACAGAGATGTTGAAAGAATCCACGGCGGTGGAA 60928901 (hg38)
(1) AAAGACAGAGATGTTGAAAGAATCC-CGGCGGTGGAA
(2) AAAGACAGAGATGTTGAAAGAATCC--GGCGGTGGAA
CSA-KO
Sequencing trace of the PCR products
(Heterozygote: 1 nucleotide deletion and 2 nucleotides deletion)
c
kDa
Ctrl
CSA-KO
72
55
36
28
17
10
CSA
H3
d
UVSSA Exon 5
sgRNA target
PAM
Ctrl Chr4:1353330 ACGAGGACAGCGACCTCGAGGAGTTTGTGCGGAGCCA 1353366 (hg38)
ACGAGGACAGCGACCTCGAGGAGTT-GTGCGGAGCCA
UVSSA-KO
(Homozygote: 1 nucleotide deletion)
e
Ctrl
UVSSA-KO
kDa
- + - + UV
250
250
55
100
15
RPB1-Ub
RPB1 S2P
CSB
CSA
UVSSA-Ub
UVSSA
H3
Chromatin

**Supplementary Fig. 1 | Schematic representation of PolII-damage PADD-seq and verification of CRISPR-Cas9 mediated gene editing, related to Fig. 1. a,** Schematic representation of PADD-seq in TCR proficient cells. Top: UV evenly induces CPDs on both template strands (TSs) and non-template strands (NTSs) independent of PolII occupancy, so the signals of PolII-associated CPDs detected by PADD-seq should be weak and symmetrically distributed on both strands right after UV irradiation. Middle: after a certain time (0.5h in our experiments), elongating PolII would be blocked by lesions on TSs but not those on NTSs, thus PADD-seq signals on TSs reflecting the level of lesion-stalled PolII should be elevated. Bottom: since *de novo* PolII release is inhibited, PADD-seq signals would decrease and finally disappear if pre-existing PolII blocked by the lesions can be resolved. **b,** Sanger sequencing trace of the CRISPR-Cas9 system edited region in *CSA*-KO cells. The sgRNA targeted region was amplified by PCR and the products were ligated to the pEASY-Blunt Zero vector. The PCR products and recombinant vectors were sequenced by Sanger sequencing. The result showed a heterozygote with 1 nucleotide deletion and 2 nucleotides deletion in exon 2 of *CSA* gene. **c,** Western blot analysis showing loss of CSA in *CSA*-KO cells. Whole cell lysates were analyzed using the indicated antibodies. H3 was used as loading control. **d,** Sanger sequencing trace of the CRISPR-Cas9 system edited region in *UVSSA*-KO cells. The sgRNA targeted region was amplified by PCR and sequenced by Sanger sequencing, showing a homozygote with 1 nucleotide deletion in exon 5 of *UVSSA* gene. **e,** Western blot analysis showing the loss of UVSSA in *UVSSA*-KO cells. The UVSSA antibody could only detect endogenous UVSSA in chromatin fraction but not in whole cell lysates due to its poor specificity, while UVSSA is recruited to chromatin and ubiquitylated after UV irradiation in the presence of CSB and CSA. Thus, cells were either irradiated with 20 J/m$^2$ UVC or not irradiated, followed by incubation for 2 h. Chromatin fractions were isolated and analyzed using the indicated antibodies. The asterisks indicate two nonspecific bands, while the lower one is close to the UVSSA-Ub band. H3 was used as loading control. Ctrl: XP-C cells. The experiments of **(c)** and **(e)** were performed once. Source data are provided as a Source Data file.

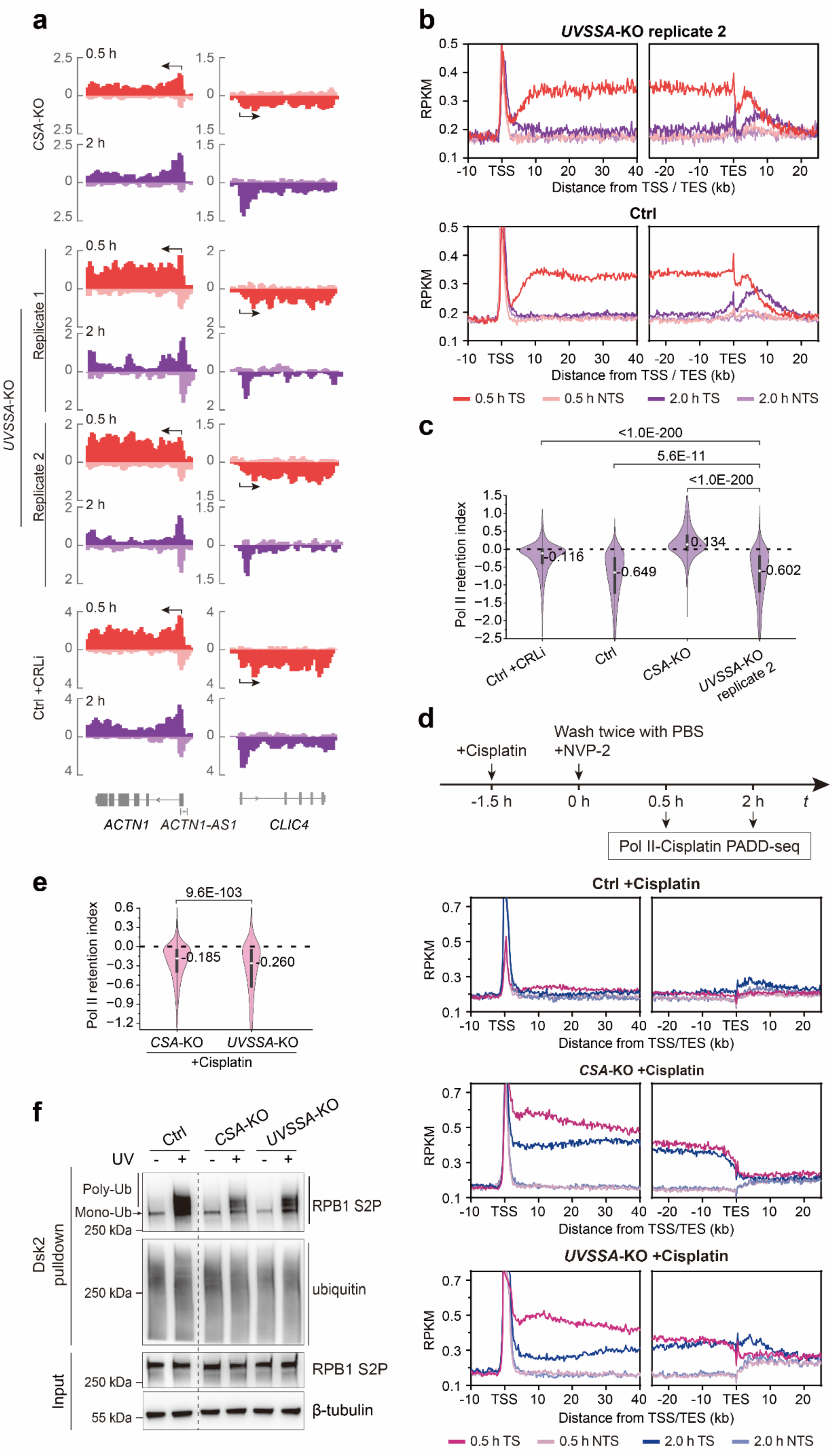
a
CSA-KO
0.5 h
2 h
UVSSA-KO
Replicate 1
Replicate 2
Ctrl +CRLi
ACTN1
ACTN1-AS1
CLIC4
b
UVSSA-KO replicate 2
Ctrl
RPKM
Distance from TSS / TES (kb)
0.5 h TS
0.5 h NTS
2.0 h TS
2.0 h NTS
c
Pol II retention index
<1.0E-200
5.6E-11
<1.0E-200
-0.116
-0.649
0.134
-0.602
Ctrl +CRLi
Ctrl
CSA-KO
UVSSA-KO replicate 2
d
+Cisplatin
Wash twice with PBS +NVP-2
-1.5 h
0 h
0.5 h
2 h
t
Pol II-Cisplatin PADD-seq
Ctrl +Cisplatin
CSA-KO +Cisplatin
UVSSA-KO +Cisplatin
Distance from TSS/TES (kb)
e
Pol II retention index
9.6E-103
-0.185
-0.260
CSA-KO
UVSSA-KO
+Cisplatin
f
Ctrl
CSA-KO
UVSSA-KO
UV
Poly-Ub
Mono-Ub
250 kDa
55 kDa
Dsk2 pulldown
Input
RPB1 S2P
ubiquitin
β-tubulin

**Supplementary Fig. 2 | Lesion-stalled PolII is resolved in *UVSSA*-KO cells by a CSA- and ubiquitination-dependent manner, related to Fig. 1. a,** Screenshots of PADD-seq results showing *ACTN1* (including its antisense gene *ACTN1-AS1*) and *CLIC4* genes under indicated conditions. **b,** Meta-gene analysis of PADD-seq signals around TSSs and TESs for active genes longer than 50 kb ($n$ = 2790) under indicated conditions. The data used in bottom panel (Ctrl) were obtained from our previous study[1]. **c,** Quantification of PolII retention on damage sites by relative change of PADD-seq signals from 0.5 h to 2 h on each gene. Active genes longer than 20 kb were selected ($n$ = 4488). $P$ value was calculated using two-tailed paired Student's $t$-test. **d,** PADD-seq for PolII-cisplatin-adduct interaction. Cells were treated with cisplatin for 1.5 h before the addition of NVP-2. Meta-gene analysis of PADD-seq signals around TSSs and TESs for active genes longer than 50 kb ($n$ = 2790) under indicated conditions. **e,** Quantification of cisplatin PADD-seq. As in **(c)** but for indicated conditions. **f,** UV-induced PolII (RPB1) ubiquitination measured by Dsk2 pulldown and western blot analysis with indicated antibodies. Cells were collected before or 30 min after UV irradiation (20 J/$m^2$). Ctrl: XP-C cells. The experiment was performed once. Source data are provided as a Source Data file.

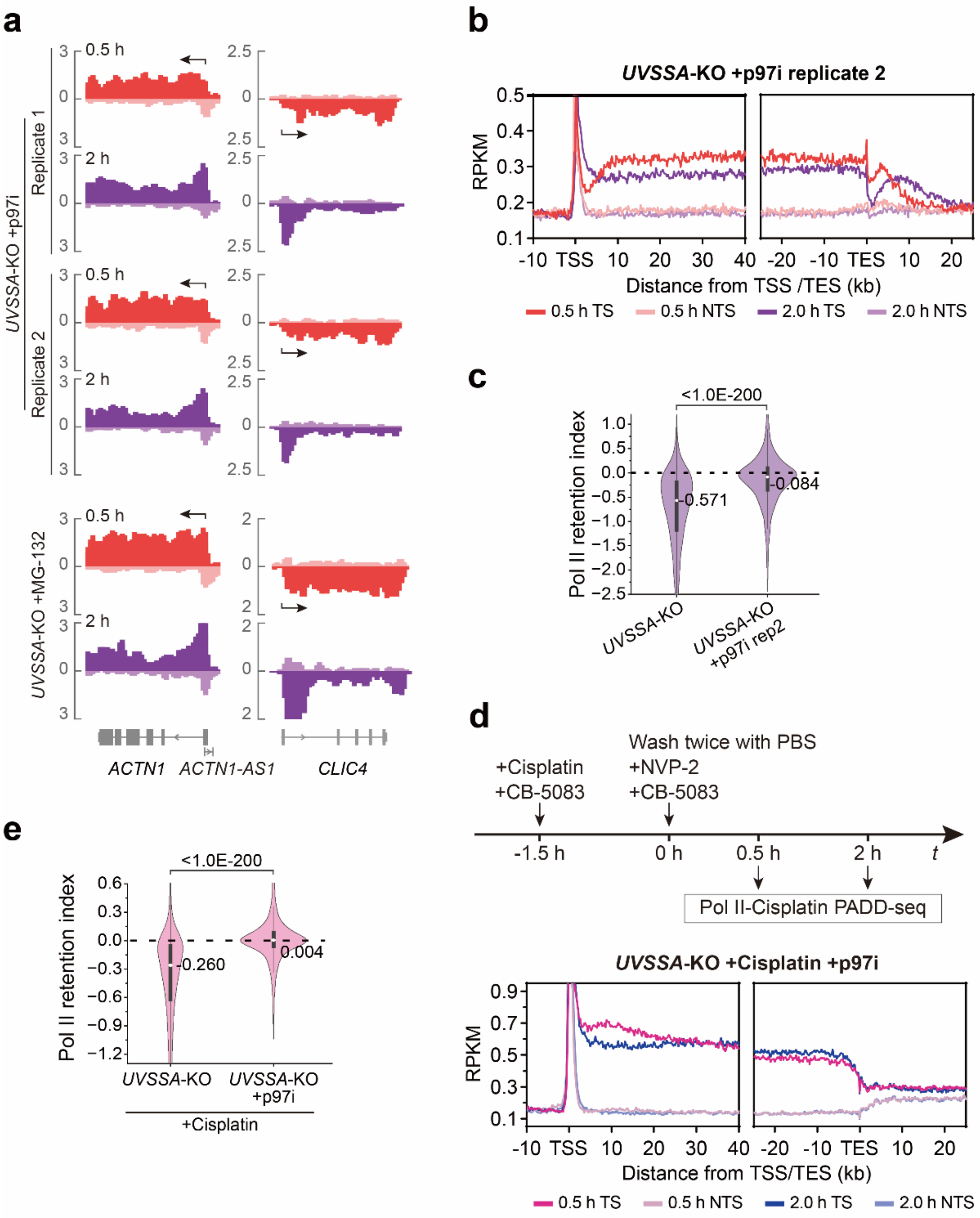
a
0.5 h
2 h
Replicate 1
Replicate 2
UVSSA-KO +p97i
UVSSA-KO +MG-132
ACTN1
ACTN1-AS1
CLIC4
b
UVSSA-KO +p97i replicate 2
RPKM
Distance from TSS /TES (kb)
0.5 h TS
0.5 h NTS
2.0 h TS
2.0 h NTS
c
<1.0E-200
Pol II retention index
-0.571
-0.084
UVSSA-KO
UVSSA-KO +p97i rep2
d
+Cisplatin
+CB-5083
Wash twice with PBS
+NVP-2
+CB-5083
-1.5 h
0 h
0.5 h
2 h
t
Pol II-Cisplatin PADD-seq
UVSSA-KO +Cisplatin +p97i
RPKM
Distance from TSS/TES (kb)
0.5 h TS
0.5 h NTS
2.0 h TS
2.0 h NTS
e
<1.0E-200
Pol II retention index
-0.260
0.004
UVSSA-KO
UVSSA-KO +p97i
+Cisplatin

**Supplementary Fig. 3 | p97 extracts PolII from damage sites in the absence of UVSSA, related to Fig. 2. a,** Screenshots of PADD-seq results showing *ACTN1* (including its antisense gene *ACTN1-AS1*) and *CLIC4* genes under indicated conditions. **b,** Meta-gene analysis of PADD-seq signals around TSSs and TESs for active genes longer than 50 kb ($n$ = 2790) under indicated conditions. **c,** Quantification of PolII retention on damage sites by relative change of PADD-seq signals from 0.5 h to 2 h on each gene. Active genes longer than 20 kb were selected ($n$ = 4488). $P$ value was calculated using two-tailed paired Student's $t$-test. **d,** PADD-seq for PolII-cisplatin-adduct interaction in *UVSSA*-KO cells treated with p97i. Experimental design was same as Supplementary Fig. 2d except CB-5083 (p97i) was added together with cisplatin and NVP-2. Meta-gene analysis of PADD-seq signals around TSSs and TESs for active genes longer than 50 kb ($n$ = 2790) under indicated conditions is shown. **e,** Quantification of cisplatin PADD-seq. As in **(c)** but for indicated conditions. Source data are provided as a Source Data file.

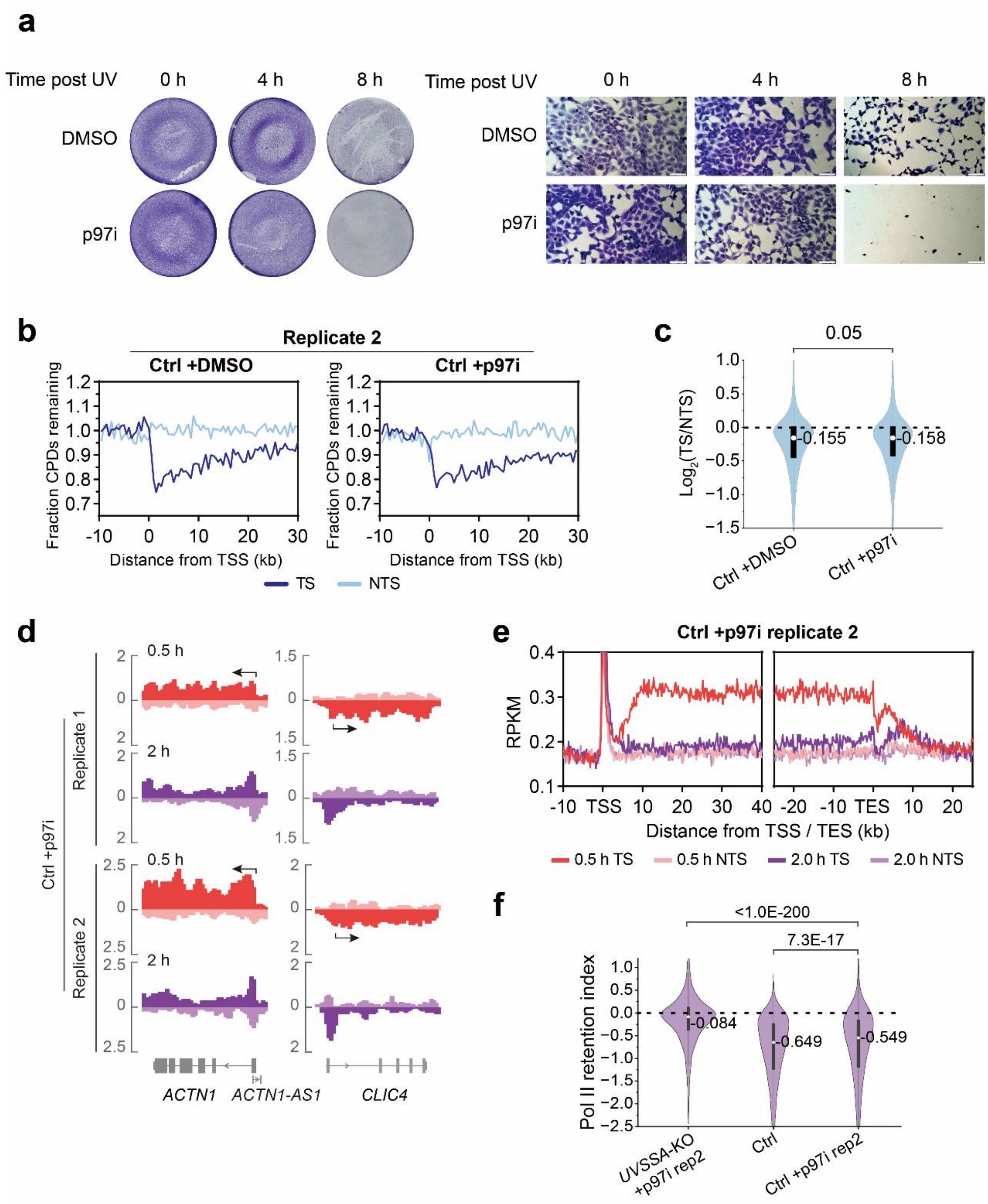
a
Time post UV
0 h
4 h
8 h
DMSO
p97i
b
Replicate 2
Ctrl +DMSO
Ctrl +p97i
Fraction CPDs remaining
Distance from TSS (kb)
TS
NTS
c
0.05
Log2(TS/NTS)
-0.155
-0.158
Ctrl +DMSO
Ctrl +p97i
d
0.5 h
2 h
Replicate 1
Replicate 2
Ctrl +p97i
ACTN1
ACTN1-AS1
CLIC4
e
Ctrl +p97i replicate 2
RPKM
Distance from TSS / TES (kb)
0.5 h TS
0.5 h NTS
2.0 h TS
2.0 h NTS
f
<1.0E-200
7.3E-17
Pol II retention index
-0.084
-0.649
-0.549
UVSSA-KO +p97i rep2
Ctrl
Ctrl +p97i rep2

**Supplementary Fig. 4 | p97 is dispensable for both TCR and PolII release in normal cells, related to Fig. 2. a,** Prolonged inhibition of p97 induces cell death. Cells were treated with 20 J/$m^2$ UV-C and 5 μM CB-5083 (or vehicle DMSO), incubated for 0 h, 4 h or 8 h, and stained with crystal violet. Pictures were taken by a normal camera (left) or a microscope (right, scale bar: 200 μm). **b,** Meta-gene analysis of Damage-seq signals around TSSs for active genes longer than 50 kb ($n$ = 2790) under indicated conditions. Cells were collected immediately (0 h) or at 4 h after UV irradiation. Fraction CPDs remaining was calculated as the ratio of 4 h to 0 h. **c,** Violin plots of relative Damage-seq signals on each active gene ($n$ = 6406). Log2 value of the ratio of fraction CPDs remaining on TS to that on NTS was calculated. **d,** Screenshots of PADD-seq results showing *ACTN1* (including its antisense gene *ACTN1-AS1*) and *CLIC4* genes under indicated conditions. **e,** Meta-gene analysis of PADD-seq signals around TSSs and TESs for active genes longer than 50 kb ($n$ = 2790) under indicated conditions. **f,** Quantification of PolII retention on damage sites by relative change of PADD-seq signals from 0.5 h to 2 h on each gene. Active genes longer than 20 kb were selected ($n$ = 4488). $P$ value was calculated using two-tailed paired Student's $t$-test for **(c)** and **(f)**. Ctrl: XP-C cells. Source data are provided as a Source Data file.

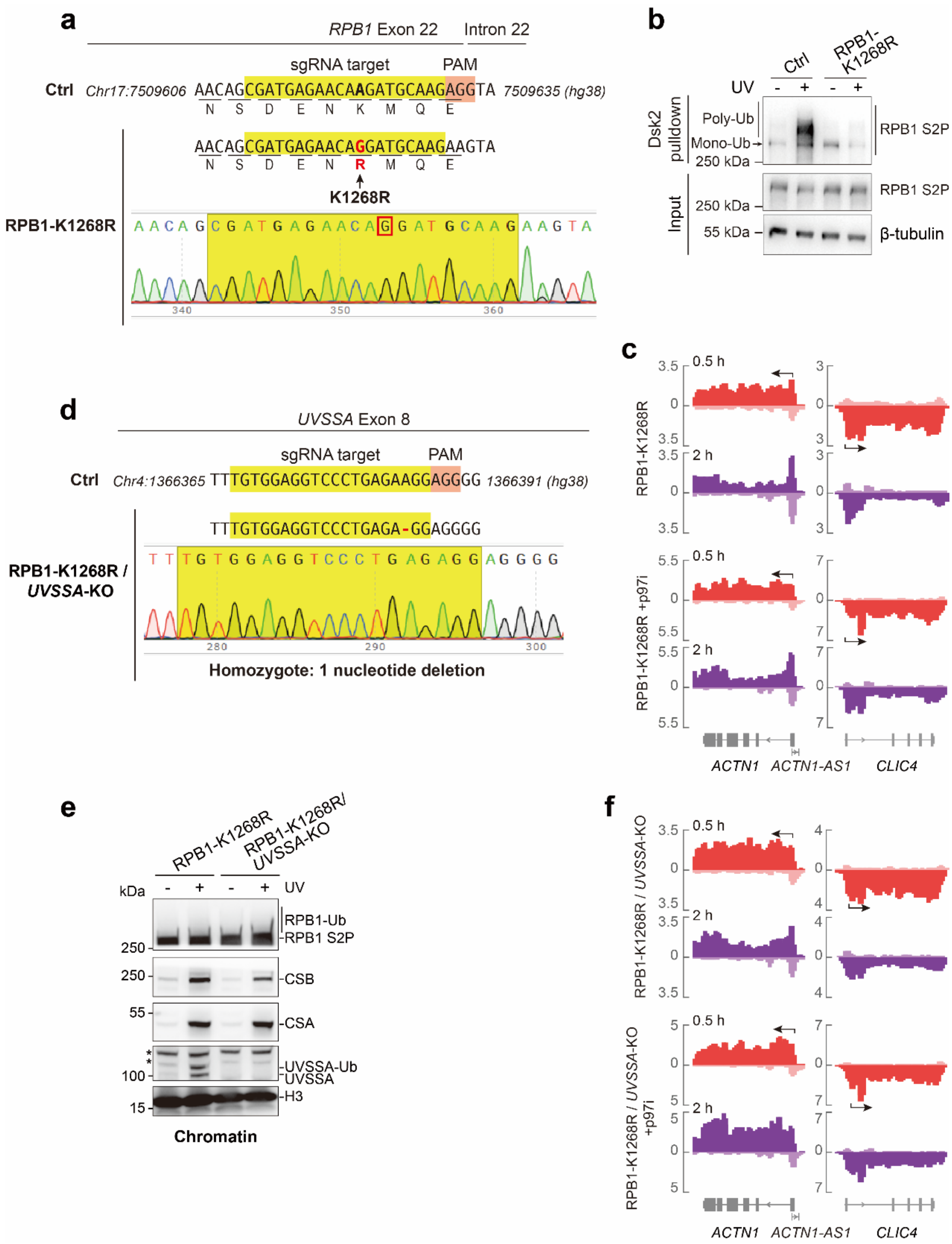
a
RPB1 Exon 22
Intron 22
sgRNA target
PAM
Ctrl
Chr17:7509606
AACAGCGATGAGAACAAGATGCAAGAGGTA
7509635 (hg38)
N S D E N K M Q E
AACAGCGATGAGAACAGGATGCAAGAAGTA
N S D E N R M Q E
K1268R
RPB1-K1268R
AACAGCGATGAGAACAGGATGCAAGAAGTA
340
350
360
b
Ctrl
RPB1-K1268R
UV
Dsk2 pulldown
Input
Poly-Ub
Mono-Ub
250 kDa
RPB1 S2P
55 kDa
β-tubulin
c
0.5 h
2 h
RPB1-K1268R
RPB1-K1268R +p97i
ACTN1
ACTN1-AS1
CLIC4
d
UVSSA Exon 8
sgRNA target
PAM
Ctrl
Chr4:1366365
TTTGTGGAGGTCCCTGAGAAGGAGGGG
1366391 (hg38)
TTTGTGGAGGTCCCTGAGA-GGAGGGG
RPB1-K1268R / UVSSA-KO
280
290
300
Homozygote: 1 nucleotide deletion
e
RPB1-K1268R
RPB1-K1268R/ UVSSA-KO
kDa
UV
RPB1-Ub
RPB1 S2P
CSB
CSA
UVSSA-Ub
UVSSA
H3
Chromatin
f
RPB1-K1268R / UVSSA-KO
RPB1-K1268R / UVSSA-KO +p97i

**Supplementary Fig. 5 | UV-induced RPB1-K1268 ubiquitination plays important but not indispensable roles in both TCR and repair-independent PolII release, related to Fig. 3. a,** Sanger sequencing trace of the CRISPR-Cas9 system edited region in RPB1-K1268R cells. The sgRNA targeted region was amplified by PCR and subjected to Sanger sequencing. The result shows a homozygous K1268R mutation of *RPB1* gene in RPB1-K1268R cells. **b,** Verification of RPB1-K1268R mutation. UV-induced RPB1 ubiquitylation was measured as in Supplementary Fig. 2f. **c,** Screenshots of PADD-seq results showing *ACTN1* (including its antisense gene *ACTN1-AS1*) and *CLIC4* genes under indicated conditions. **d,** Sanger sequencing trace of the CRISPR-Cas9 system edited region in exon 8 of *UVSSA* gene in RPB1-K1268R/*UVSSA*-KO double mutant cells. The sgRNA targeted region was amplified by PCR and subjected to Sanger sequencing. The result showed a homozygous 1 nucleotide deletion of *UVSSA* gene in RPB1-K1268R/*UVSSA*-KO cells. **e,** Western blot analysis showing the loss of UVSSA in RPB1-K1268R/*UVSSA*-KO cells as in Supplementary Fig. 1e. **f,** As in **(c)** but for indicated conditions. Ctrl: XP-C cells. The experiments of **(b)** and **(e)** were performed once. Source data are provided as a Source Data file.

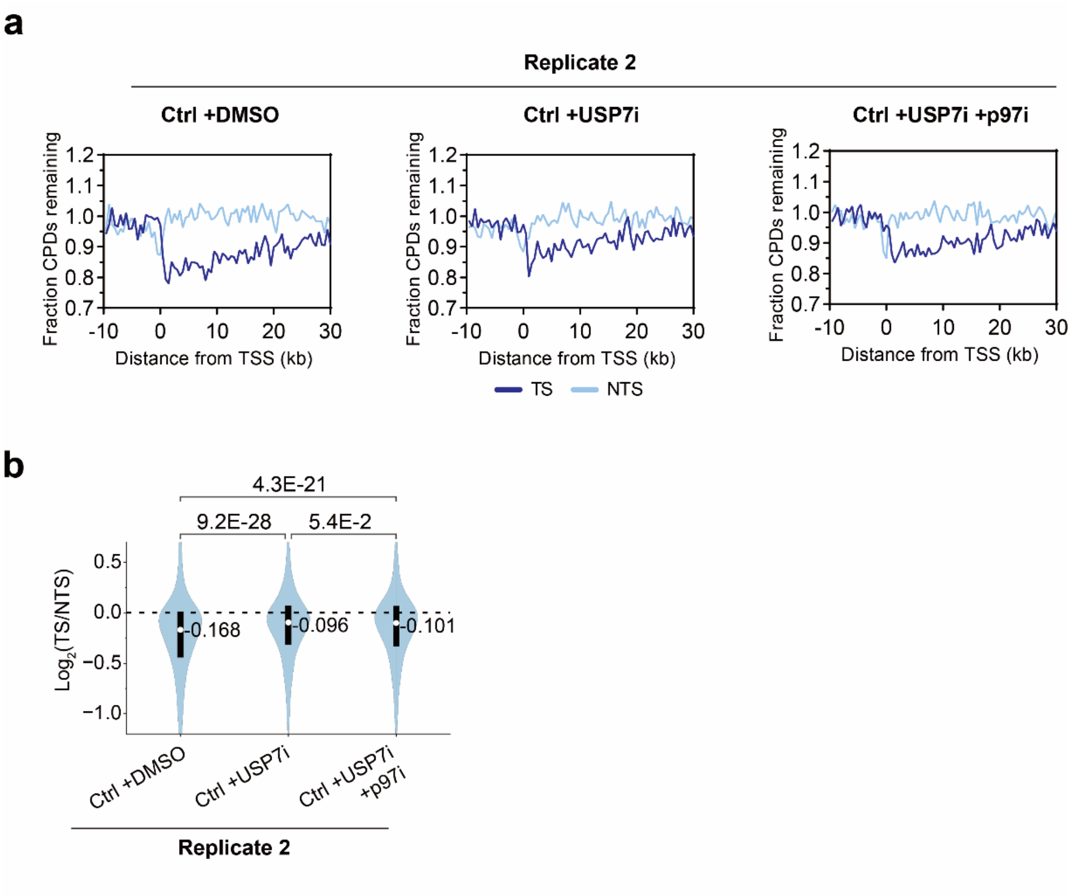


**Supplementary Fig. 6. USP7 is involved in TCR, related to Fig. 4. a,** Meta-gene analysis of Damage-seq signals around TSSs for active genes longer than 50 kb ($n$ = 2790) under indicated conditions. Cells were collected immediately (0 h) or at 4 h after UV irradiation. Fraction CPDs remaining was calculated as the ratio of 4 h to 0 h. **b,** Violin plots of relative Damage-seq signals on each active gene ($n$ = 6406). Log2 value of the ratio of fraction CPDs remaining on TS to that on NTS was calculated. $P$ value was calculated using two-tailed paired Student's $t$-test. Source data are provided as a Source Data file.

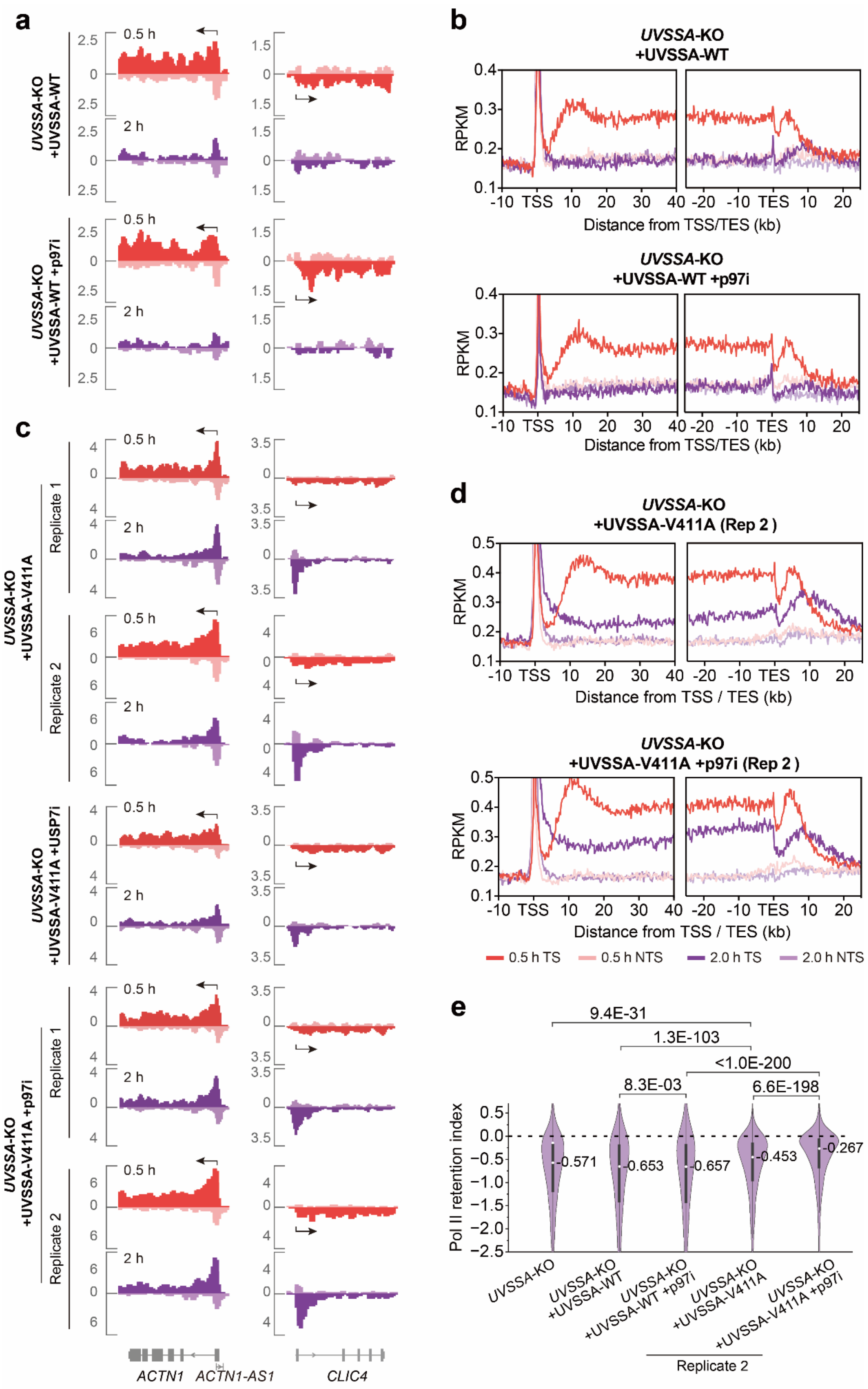
a
UVSSA-KO +UVSSA-WT
UVSSA-KO +UVSSA-WT +p97i
0.5 h
2 h
b
UVSSA-KO +UVSSA-WT
UVSSA-KO +UVSSA-WT +p97i
RPKM
Distance from TSS/TES (kb)
c
UVSSA-KO +UVSSA-V411A
UVSSA-KO +UVSSA-V411A +USP7i
UVSSA-KO +UVSSA-V411A +p97i
Replicate 1
Replicate 2
ACTN1
ACTN1-AS1
CLIC4
d
UVSSA-KO +UVSSA-V411A (Rep 2 )
UVSSA-KO +UVSSA-V411A +p97i (Rep 2 )
Distance from TSS / TES (kb)
0.5 h TS
0.5 h NTS
2.0 h TS
2.0 h NTS
e
9.4E-31
1.3E-103
<1.0E-200
8.3E-03
6.6E-198
Pol II retention index
-0.571
-0.653
-0.657
-0.453
-0.267
UVSSA-KO
UVSSA-KO +UVSSA-WT
UVSSA-KO +UVSSA-WT +p97i
UVSSA-KO +UVSSA-V411A
UVSSA-KO +UVSSA-V411A +p97i
Replicate 2

**Supplementary Fig. 7 | USP7 cannot abolish repair-independent PolII release driven by p97, related to Fig. 5. a,** Screenshots of PADD-seq results showing *ACTN1* (including its antisense gene *ACTN1-AS1*) and *CLIC4* genes under indicated conditions. **b,** Meta-gene analysis of PADD-seq signals around TSSs and TESs for active genes longer than 50 kb ($n$ = 2790) under indicated conditions. **c,** As in **(a)** but for indicated conditions. **d,** As in **(b)** but for indicated conditions. **e,** Quantification of PolII retention on damage sites by relative change of PADD-seq signals from 0.5 h to 2 h on each gene. Active genes longer than 20 kb were selected ($n$ = 4488). $P$ value was calculated using two-tailed paired Student's $t$-test. Source data are provided as a Source Data file.

**Supplementary Table 1. Oligonucleotides sequences used for gene editing by CRISPR-Cas9**

| Target gene | sgRNA sequence (5’-3’) | Homology directed repair (HDR) oligos | Description |
|---|---|---|---|
| *CSA* | GATGTTGAAAG AATCCACGG | N/A | Knock out *CSA* in XP-C cells |
| *RPB1* | CGATGAGAACA AGATGCAAG | AGGTTTTGGTGACGACT TGAACTGCATCTTTAAT GATGACAATGCAGAGA AGCTGGTGCTCCGTATT CGCATCATGAACAGCGA TGAGAACAGGATGCAA GAGGTAATGGGGGTCCT AGAAGTCAGCGTG | Generate RPB1-K1268R mutated XP-C cells |
| *UVSSA* | GCGACCTCGAG GAGTTTGTG | N/A | Knock out *UVSSA* in XP-C cells |
| *UVSSA* | TGTGGAGGTCC CTGAGAAGG | N/A | Knock out *UVSSA* in RPB1-K1268R mutated XP-C cells |

**Supplementary Table 2. Primers used for site directed mutagenesis to generate UVSSA-V411A mutant**

| Primer | Sequences (5’-3’) |
|---|---|
| Forward primer | ACGATGAGGACTTTGTGGAGGCCCCTGAGAAGGAGGGGTATGA |
| Reverse primer | TCATACCCCTCCTTCTCAGGGGCCTCCACAAAGTCCTCATCGT |

**Supplementary Table 3. Oligonucleotides sequences used for Damage-seq and PADD-seq**

| Oligonucleotides | Sequences (5’-3’) |
|---|---|
| Ad1T | /5Phos/GATCGGAAGAGCACACGTCTGAACTCCAGTCA/3SpC3/ |
| Ad1B | NNNNNGACTGGTTCCAATTGAAAGTGCTCTTCCGATC*T |
| O3P | GACTGGAGTTCAGACGTGTGCTCTTCCGATCT |
| Ad2T | /5Phos/AGATCGGAAGAGCGTCGTGTAGGGAAAGAGTGT/3SpC3/ |
| Ad2B | ACACTCTTTCCCTACACGACGCTCTTCCGATCTNNNNNN/3SpC3/ |

The “/5Phos/” indicates 5’ phosphorylation; “/3SpC3/” indicates 3’ C3-spacer modification which blocks the 3’ end; “*” indicates phosphorothioate bond. Oligonucleotides were purchased from Sangon Biotech. Ad1 was prepared by annealing Ad1T and Ad1B. Ad2 was prepared by annealing Ad2T and Ad2B. For annealing, 100 µM of each oligo was mixed in 50 µl of hybridization buffer (0.5 mM Tris-HCl pH 8.0, 100 mM NaCl, and 0.05 mM EDTA), boiled for 2 min, naturally cooled down to room temperature, and stored at -20 °C.
